# Spatially and Functionally Distinct mTORC1 Entities Orchestrate the Cellular Response to Amino Acid Availability

**DOI:** 10.1101/2023.10.03.559930

**Authors:** Stephanie A. Fernandes, Danai-Dimitra Angelidaki, Julian Nüchel, Jiyoung Pan, Peter Gollwitzer, Yoav Elkis, Filippo Artoni, Sabine Wilhelm, Marija Kovacevic-Sarmiento, Constantinos Demetriades

## Abstract

Amino acid (AA) availability is a robust determinant of cell growth, through controlling mTORC1 activity ^1^. According to the predominant model in the field, AA sufficiency drives the recruitment and activation of mTORC1 on the lysosomal surface by the heterodimeric Rag GTPases, from where it coordinates the majority of cellular processes (reviewed in ^2,3^). Importantly, however, 15 years after its initial discovery, the teleonomy of the proposed lysosomal regulation of mTORC1, and where mTORC1 acts on its effector proteins remain enigmatic ^4^. Here, by using multiple pharmacological and genetic means to perturb the lysosomal AA sensing and protein recycling machineries, we describe the spatial separation of mTORC1 regulation and downstream functions in mammalian cells, with lysosomal and non-lysosomal mTORC1 phosphorylating distinct substrates in response to different AA sources. Moreover, we reveal that a fraction of mTOR localizes at lysosomes due to basal lysosomal proteolysis that locally supplies new AAs, even in cells grown in the presence of extracellular nutrients, whereas cytoplasmic mTORC1 is regulated by exogenous AAs. Overall, our study substantially expands our knowledge about the topology of mTORC1 regulation by AAs, and hints at the existence of distinct, Rag- and lysosome-independent mechanisms that control its activity at other subcellular locations. Given the importance of mTORC1 signalling and AA sensing for human ageing and disease ^2^, our findings will likely open new directions toward the identification of function-specific mTORC1 regulators, and suggest new targets for drug discovery against conditions with dysregulated mTORC1 activity in the future.

## Introduction

Cell growth is a crucial and tightly regulated process. Cells take up nutrients, like AAs, from their environment and use them to synthesize complex macromolecules, which they incorporate to increase their mass and grow. As growth is very energy consuming, cells have developed mechanisms to sense nutrient availability and to adjust their metabolism accordingly, so that they only grow when conditions are optimal. These mechanisms are of great importance, as dysregulation of growth can be detrimental for cellular and organismal health ^5–7^.

The mechanistic/mammalian Target of Rapamycin Complex 1 (mTORC1) is a master regulator of cellular growth and metabolism. It functions as a sensor and a molecular rheostat that links information from the cellular milieu to the physiological and metabolic properties of the cells ^5,8–11^. The availability of AAs, in particular, is one of the most powerful signals for mTORC1 activation. In fact, AA signalling can override other stimuli, e.g., growth factor availability ^1,12–14^. Because AAs are the building blocks to make proteins, and mTORC1 controls protein synthesis, this mechanism ensures that cells produce proteins only when AAs are available. In addition to protein synthesis, mTORC1 activity affects the majority of cellular functions, and, as a result, it can influence organismal health, lifespan and ageing. Hyperactivation of mTORC1—caused mainly by mutations in its upstream regulators—is of clinical relevance and a common feature of most cancer types and several metabolic disorders ^8,15–17^. Moreover, AA signalling to mTORC1 is medically relevant too, as nutritional AA overload has been linked to obesity and diabetes via hyperactivation of mTORC1 ^18^. Therefore, how AAs regulate mTORC1 is an important question, relevant for both basic and clinical research.

Work over the last 15 years has built a lysosome-centric model of mTORC1 regulation by AAs, based on which, AAs control the subcellular localization of mTORC1 via regulating the activity of the lysosomal heterodimeric Rag GTPases, composed of RagA or RagB bound to RagC or RagD. Under conditions of AA sufficiency, an ‘active’ Rag dimer (GTP-bound RagA/B and GDP-bound RagC/D) recruits mTORC1 to the lysosomal surface where it is activated by another small GTPase called Rheb ^19–22^. In contrast, AA starvation leads to inactivation of the Rags (GDP-bound RagA/B and GTP-bound RagC/D) and the subsequent release of mTORC1 from the lysosomal surface, which has been linked to an impairment of its activity ^19,23,24^. Following the original discovery of the Rag GTPases as a central hub in AA sensing ^19,20^, the quest to understand how cells sense the availability of AAs over the following years has led to the identification of a large number of lysosomal and cytoplasmic proteins that modify Rag dimer activity in an AA-dependent manner (reviewed in ^2,25,26^).

According to this model, mTORC1 is exclusively activated by AAs on the lysosomal surface, from where it controls all cellular processes, while dissociation of mTORC1 from lysosomes is linked to its inactivation. However, scattered evidence in the literature suggests that the cell biology of AA signalling to mTORC1 likely is more complex than currently appreciated. Indeed, although a fraction of mTORC1 does localize to lysosomes in cells grown under AA-replete conditions, a substantial amount of this complex is found away from lysosomes ^27,28^. Whether this non-lysosomal mTORC1 pool is active or inactive is an important open question. Furthermore, many of the canonical mTORC1 effectors are known to be non-lysosomal proteins, hence where and how mTORC1 acts on its various substrates is, so far, unclear. Finally, because mTORC1 activity (as assessed by the phosphorylation of its most-commonly-used substrate, S6K) acutely responds to changes in exogenous AA sufficiency, the teleonomy of the exclusively lysosomal regulation of mTORC1 by AAs remains enigmatic. Taken together, a model where mTORC1 is exclusively present and regulated on the lysosomal surface appears counterintuitive.

## Results

### Basal lysosomal proteolysis and local AA production is responsible for the lysosomal recruitment and regulation of mTORC1

Previous efforts to study how AAs signal to regulate mTORC1 have focused on the lysosomally anchored Rag GTPases and revealed a complex network of proteins that regulate Rag activity and function ^2,25,26^. However, the reason for the lysosomal localization of mTORC1 in cells grown in the presence of AAs has been a long-standing question in the field. Given that lysosomes are the main degradative organelles in cells, we intuitively hypothesised that local AA production, due to basal protein recycling inside lysosomes, may be playing an important role in this process. Bafilomycin A1 (BafA1) is a macrolide antibiotic that blocks lysosomal function by targeting the V-ATPase (vacuolar H^+^ ATPase), causing alkalinization of the lysosomal lumen and preventing protease activity, AA efflux, and autolysosome formation ^21,29^ (Fig. 1a). Consistent with previous reports ^30^, treatment of human embryonic kidney HEK293FT cells with BafA1 caused strong accumulation of LC3B (Fig. 1b,c and Ext. Data Fig. 1a) and increased the levels of the macro- and micro-autophagy adaptor proteins TAX1BP1, NBR1 (Ext. Data Fig. 1a), and p62 (Ext. Data Fig. 1b,c), confirming that basal lysosomal proteolysis is robustly active in these cells, even when they are grown in the presence of exogenous AAs. Similar results were obtained in mouse embryonic fibroblasts (MEFs) (Ext. Data Fig. 1d,e), showing that this phenomenon is not cell-type-or species-specific.

**Figure 1.**
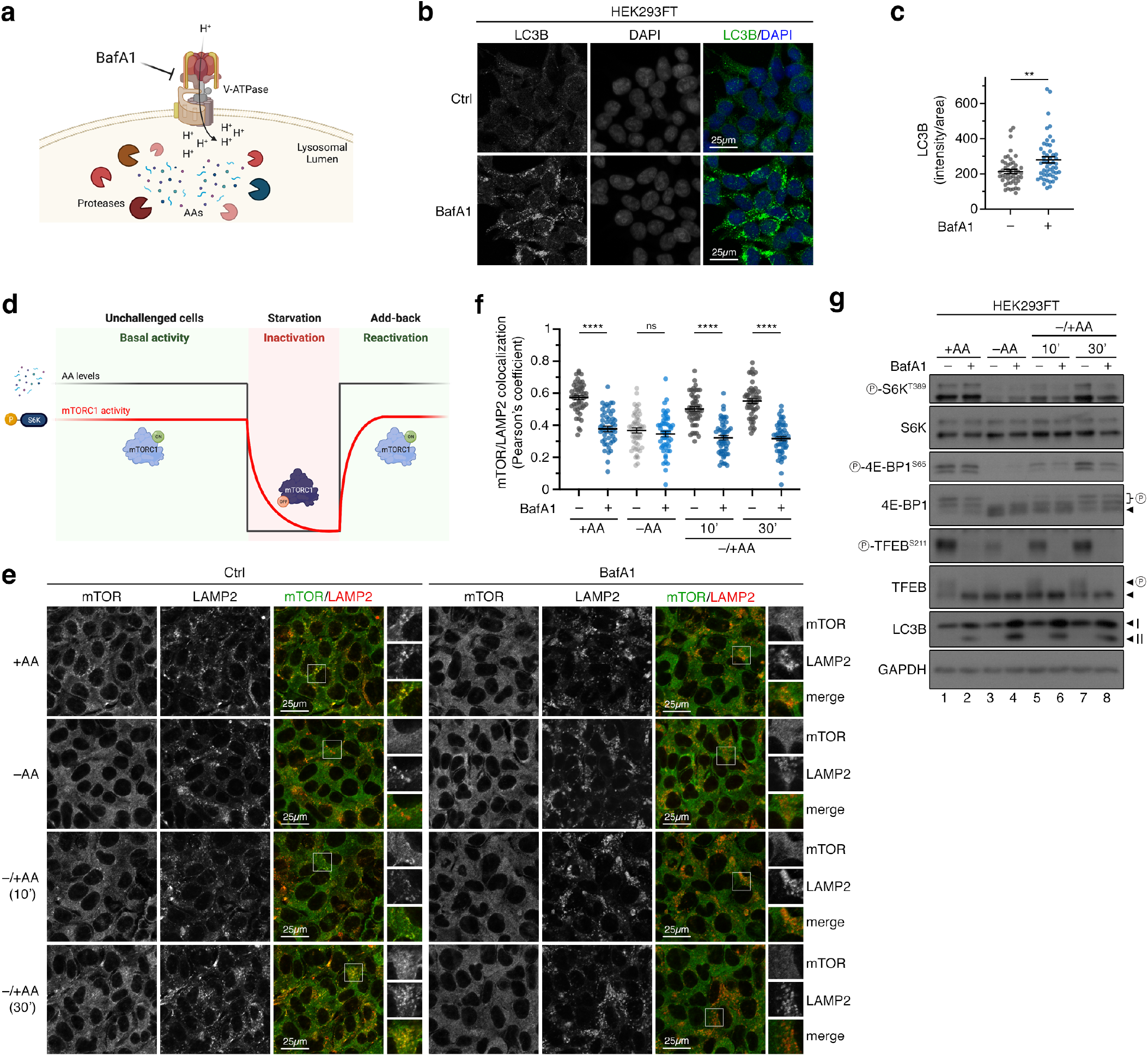
Blockage of lysosomal function disconnects mTORC1 localization and substrate-specific activity. **(a)** Schematic model of pharmacological inhibition of lysosomal function by Bafilomycin A1 (BafA1) targeting the V-ATPase. **(b-c)** Basal lysosomal proteolysis in HEK293FT cells shown by accumulation of LC3B upon BafA1 treatment (b). Quantification of LC3B signal in (c). n = 50 individual cells from 5 independent fields per condition. **(d)** Schematic representation of the treatment strategy followed in this manuscript, assessing mTORC1 activity under basal (unchallenged cells), starvation, or acute re-activation (AA add-back) conditions. AA levels shown by black line, mTORC1 activity shown by red line. See methods for a detailed description. **(e-f)** Colocalization analysis of mTOR with LAMP2 (lysosomal marker) in HEK293FT WT cells, treated as indicated in the figure, using confocal microscopy. Scale bars = 25 μm. Magnified insets shown to the right (e). Quantification of colocalization in (f). n = 50 individual cells from 5 independent fields per condition. **(g)** Immunoblots with lysates from HEK293FT WT cells, treated with media containing or lacking AAs, in basal (+AA), starvation (–AA) or add-back (–/+AA) conditions, and BafA1 as shown, probed with the indicated antibodies. Arrowheads indicate bands corresponding to different protein forms, when multiple bands are present. P: phosphorylated form. Data shown are representative of 3 replicate experiments. Data in (c), (f) shown as mean ± SEM. ** p<0.01, **** p<0.0001, ns: non-significant. See also Ext. Data Figures 1-2.

mTORC1 localization and activity respond acutely and dynamically to changes in extracellular AA availability (Fig. 1d-g). Notably, although mTORC1 can be active and is found on lysosomes under either basal or AA add-back conditions (Fig. 1d-g), our previous work suggested that the mechanistic details of its activation (as indicated by the phosphorylation of its best-described substrate, S6K) may differ depending on the specific treatment strategy (see Fig. S1D in ^24^). Most importantly, the involvement of virtually all components of the lysosomal AA sensing machinery, including the Rags—as well as the upstream network that signals AA sufficiency via the Rags—has been studied using very consistently an ‘add-back’ treatment strategy, originally introduced by the Sabatini lab (50 minutes of AA starvation, followed by 10 minutes of AA re-addition). Apparently, this protocol only studies the acute re-activation of mTORC1, while overlooking the basal mTORC1 activation in unchallenged cells. Therefore, for subsequent experiments, we studied the regulation of mTORC1 under all aforementioned nutritional conditions (basal, starvation, add-back of AAs) (Fig. 1d). In control HEK293FT cells, mTOR showed a mixed localization pattern with part of the signal colocalizing with LAMP2 (used as a lysosomal marker) and part being diffusely cytoplasmic. As opposed to control cells, mTOR no longer colocalized with lysosomes in cells treated with BafA1, regardless of the treatment regimen (Fig. 1e,f), thereby supporting our initial hypothesis about basal protein recycling and local AA production in lysosomes being the cause for the lysosomal recruitment of mTORC1. Next, we assessed the effect of BafA1 on the dynamics of mTORC1 activation by AAs. Unlike most previous studies that focused on specific substrates like S6K to assay mTORC1 activity, we here tested multiple mTORC1 substrates. Interestingly, these experiments showed that phosphorylation of S6K and 4E-BP1, two canonical substrates of mTORC1, was largely unaffected by BafA1 under basal conditions (Fig. 1g). In contrast, the re-activation of mTORC1 upon AA re-addition was compromised in BafA1-treated cells (Fig. 1g), in line with a previous report using concanamycin A (ConA) to block lysosomal function and AA efflux from these organelles ^27^. Strikingly, unlike for S6K and 4E-BP1, the phosphorylation of TFEB, a lysosomal non-canonical mTORC1 substrate, was abolished by BafA1 under all conditions tested (Fig. 1g), paralleling the loss of lysosomal mTORC1 accumulation (Fig. 1e,f). Of note, culturing cells in starvation media specifically lacking AAs readily caused the dephosphorylation of all mTORC1 substrates in both control and BafA1-treated cells (Fig. 1g). Similar results were obtained by i) using concanamycin A (ConA), an independent v-ATPase inhibitor (Ext. Data Fig. 2a-c); ii) using chloroquine (CQ), a lysosomotropic weak base that alkalinizes the lysosomal lumen independently of v-ATPase inhibition (Ext. Data Fig. 2d-f); iii) specifically inhibiting the activity of lysosomal proteases with a combination of pepstatin A (PepA) and E64 that does not affect lysosomal acidification (Fig. 2a-d); or iv) by preventing the delivery of lysosomal enzymes to this compartment via transient knockdown or knockout (KO) of GNPTAB (N-acetylglucosamine-1-phosphotransferase subunits alpha/beta), a protein responsible for proper lysosomal enzyme sorting at the Golgi (Fig. 3a-h).

**Figure 2.**
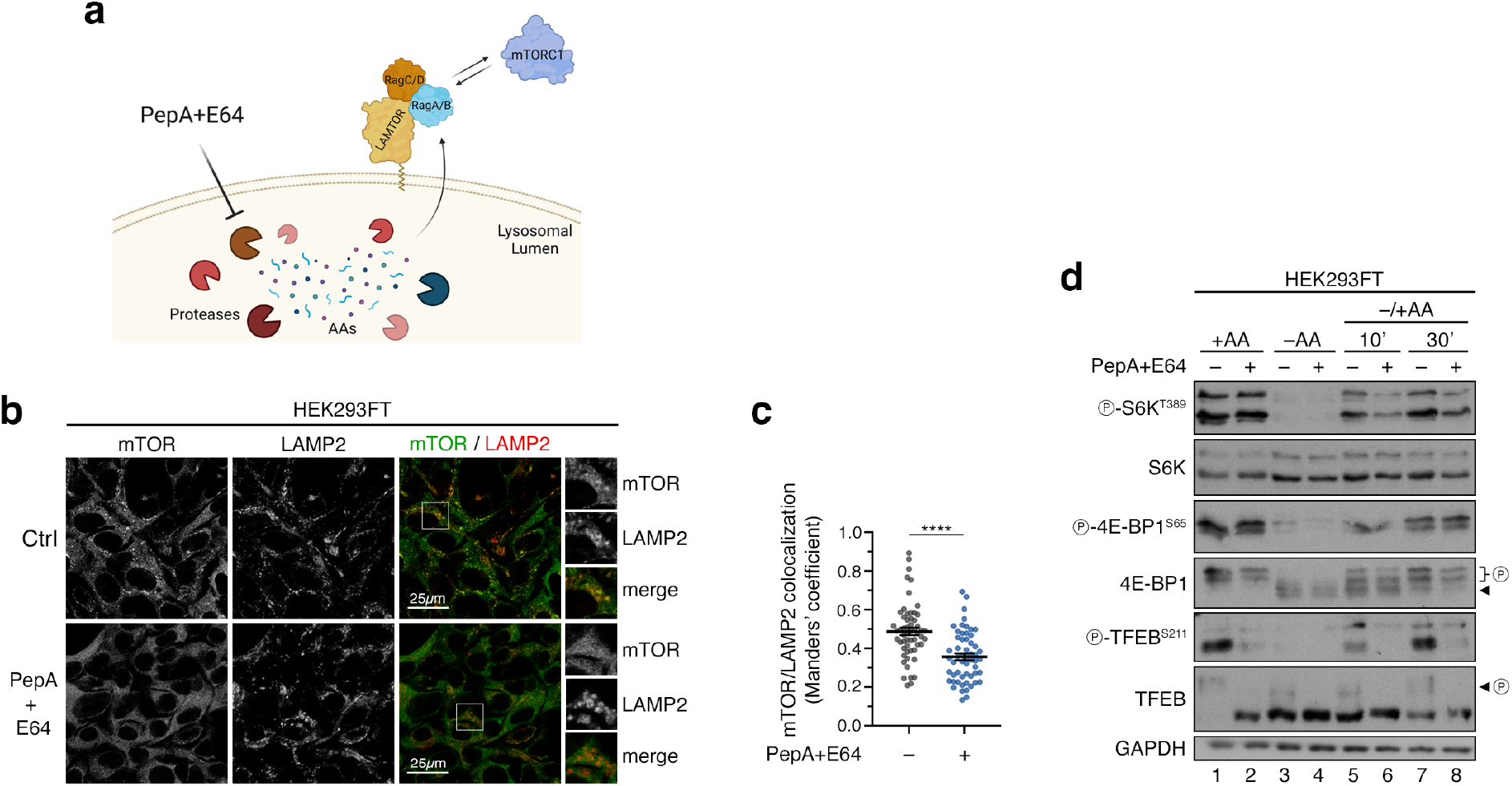
Blockage of lysosomal protease activity disconnects mTORC1 localization on lysosomes and activity toward cytoplasmic substrates. **(a)** Schematic model of pharmacological inhibition of lysosomal proteases by pepstatin A (PepA) and E64 blocking local AA production. **(b-c)** Colocalization analysis of mTOR with LAMP2 (lysosomal marker) in HEK293FT WT cells, treated as indicated in the figure, using confocal microscopy. Scale bars = 25 μm. Magnified insets shown to the right (b). Quantification of colocalization in (c). n = 56 individual cells from 3 independent fields per condition. **(d)** Immunoblots with lysates from HEK293FT WT cells, treated with media containing or lacking AA, in basal (+AA), starvation (–AA) or add-back (–/+AA) conditions, and protease inhibitors (PepA+E64) as shown, probed with the indicated antibodies. Arrowheads indicate bands corresponding to different protein forms, when multiple bands are present. P: phosphorylated form. Data in (c) shown as mean ± SEM. **** p<0.0001.

**Figure 3.**
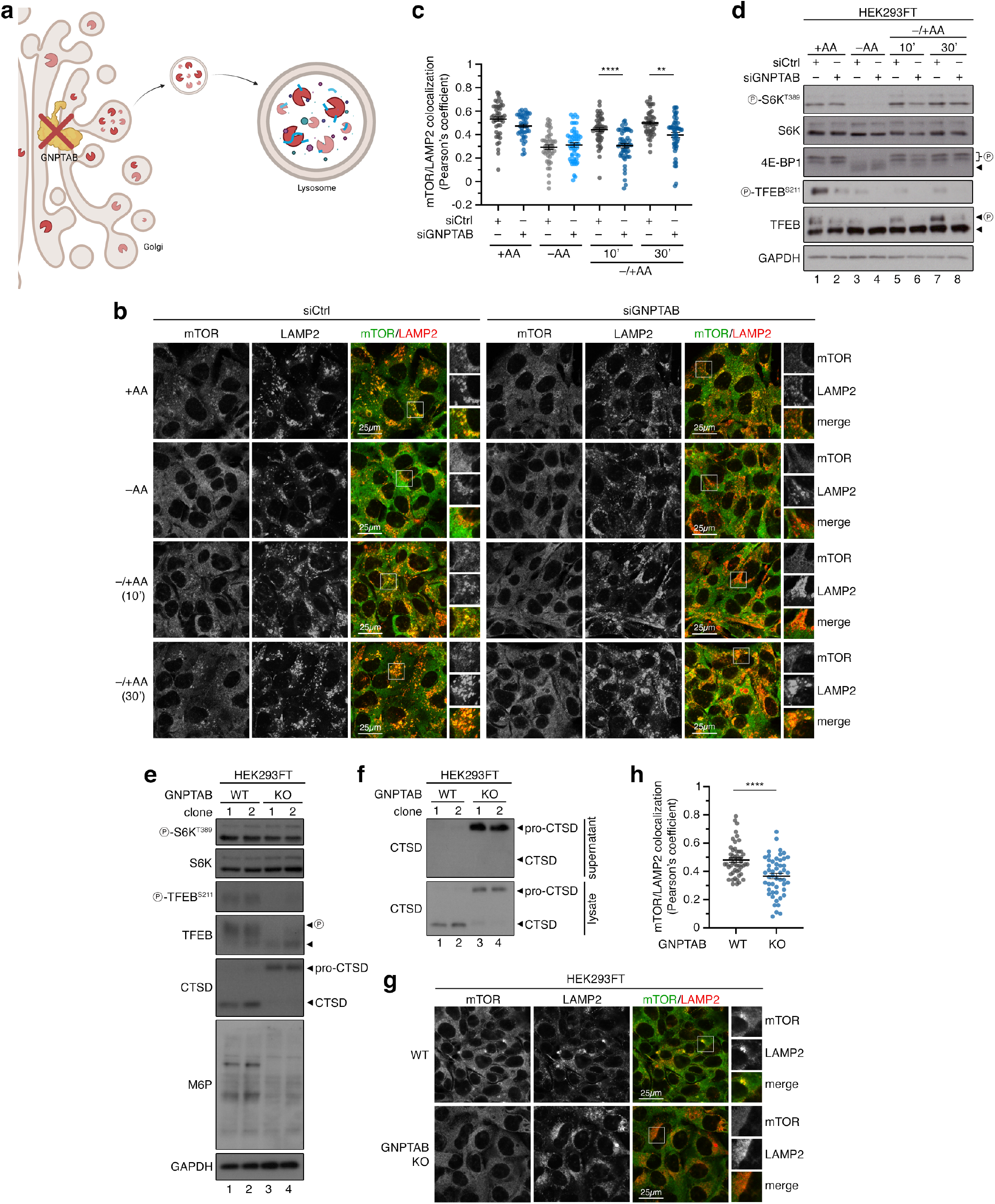
Blockage of proper lysosomal enzyme sorting and delivery disconnects mTORC1 localization on lysosomes and activity toward cytoplasmic substrates. **(a)** Schematic model of lysosomal enzyme sorting at the Golgi and delivery to lysosomes that depends on the GNPTAB enzyme. **(b-c)** Colocalization analysis of mTOR with LAMP2 (lysosomal marker) in HEK293FT WT or RagA/B KO cells using confocal microscopy. Cells were transiently transfected with siRNAs targeting GNPTAB or a control RNAi duplex (siCtrl) and treated as indicated in the figure. Scale bars = 25 μm. Magnified insets shown to the right (b). Quantification of mTOR/LAMP2 colocalization in (c). n = 44-50 individual cells from 5 independent fields per condition. Representative data from one out of three independent experiments are shown. **(d)** Immunoblots with lysates from HEK293FT WT cells, transiently transfected with siRNAs targeting GNPTAB or a control RNAi duplex (siCtrl), and treated with media containing or lacking AA, in basal (+AA), starvation (–AA) or add-back (–/+AA) conditions, probed with the indicated antibodies. n = 3 independent experiments. **(e-f)** Functional characterization of GNPTAB KO HEK293FT cells. Two independent knockout clones show impaired cathepsin D (CTSD) processing and mannose-6-phosphate (M6P)-tagging of proteins. Note also the differential effect of GNPTAB loss on the different mTORC1 substrates (TFEB de-phosphorylated in KO cells, whereas S6K phosphorylation is largely unaffected) (e). The pro-form of CTSD is aberrantly secreted in the medium of GNPTAB KO cells (f). n = 3 independent experiments. **(g-h)** Lysosomal accumulations of mTOR are lost in GNPTAB KOs (g). Quantification of mTOR/LAMP2 colocalization in (h). n = 50 individual cells from 5 independent fields per condition. Representative data from one out of three independent experiments are shown. Arrowheads indicate bands corresponding to different protein forms, when multiple bands are present. P: phosphorylated form. Data in graphs shown as mean ± SEM. ** p<0.01, **** p<0.0001.

To test if the persistent phosphorylation of S6K and 4E-BP1 in BafA1-treated cells is due to sustained activity of non-lysosomal mTORC1 or due to slower dephosphorylation kinetics, we performed BafA1 time-course experiments looking both at mTOR localization and activity, using S6K, 4E-BP1, and TFEB phosphorylation as read-outs. These experiments showed that mTOR lysosomal accumulations disappear already after 2 hours of BafA1 treatment (Fig. 4a,b), closely correlating with the kinetics of TFEB dephosphorylation (Fig. 4c,d). In stark contrast, phosphorylation of S6K and 4E-BP1 stayed largely unaffected for the whole duration of the experiment (Fig. 4c,e). Dephosphorylation kinetics typically depend on the counteracting activities of a kinase and the respective phosphatase. Accordingly, by blocking mTORC1 kinase activity in control and RagA/B KO cells (in which mTOR is no longer lysosomal ^19,23,24^, like in BafA1-treated cells; see also below) with rapamycin for different times, we looked at the rate of S6K dephosphorylation as a means to assess phosphatase activity. Notably, these experiments showed that phosphatase activity toward S6K is comparable between the two genotypes (Fig. 4f,g), further supporting that the persistent phosphorylation of cytoplasmic mTORC1 substrates in cells with non-lysosomal mTORC1 is not due to slower or impaired dephosphorylation (i.e., due to compromised phosphatase activity).

**Figure 4.**
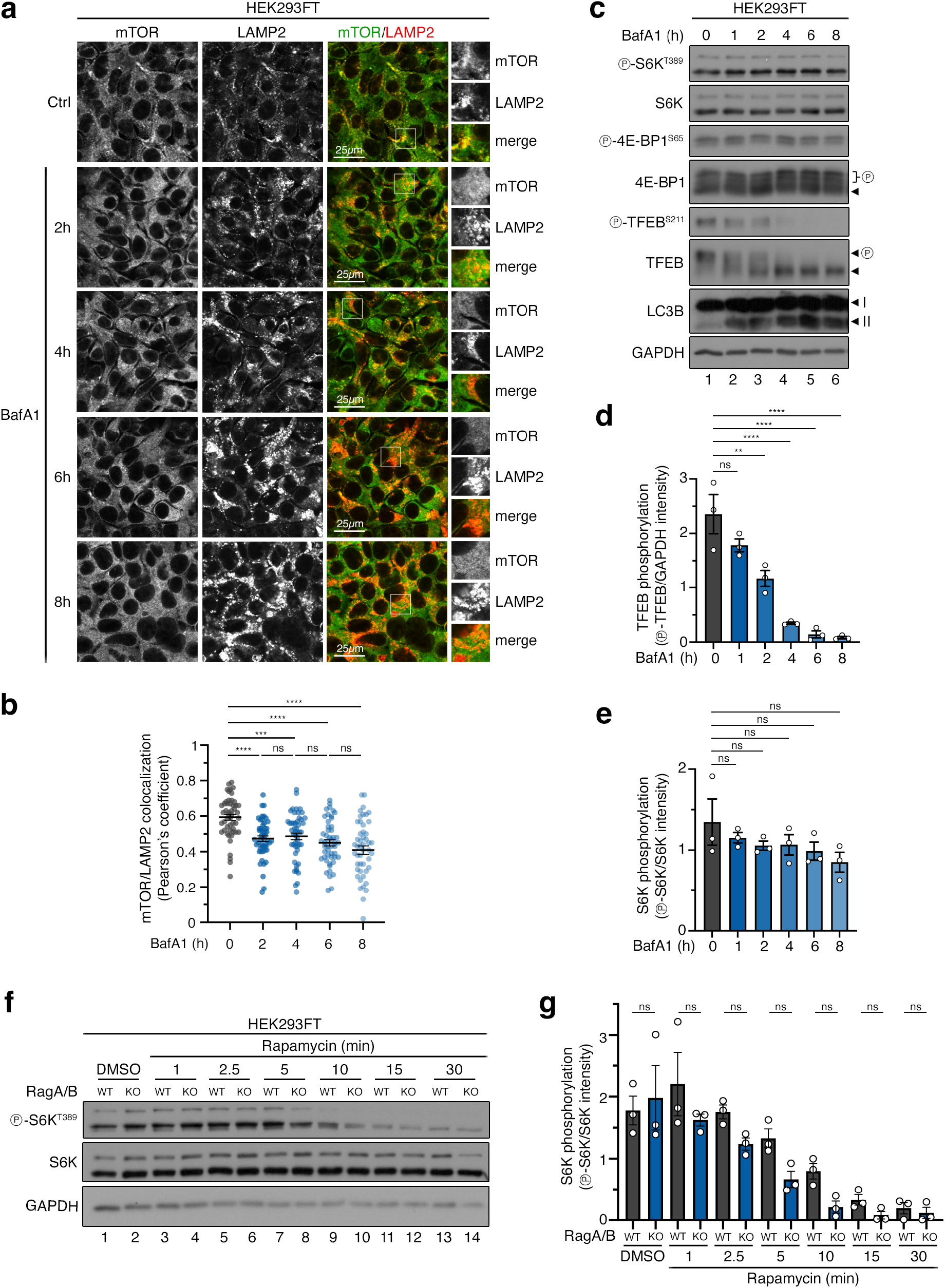
The persistent phosphorylation of cytoplasmic mTORC1 substrates in cells with non-lysosomal mTORC1 is not due to impaired or slower substrate dephosphorylation. **(a-b)** mTOR delocalizes away from lysosomes already after 2h of BafA1 treatment. Time course of BafA1 treatment (2-8 h) to block lysosomal function (a). Quantification of mTOR/LAMP2 colocalization in (b). n = 50 individual cells from 5 independent fields per condition. Representative data from one out of three independent experiments are shown. **(c-e)** Dephosphorylation kinetics of lysosomal (TFEB) and cytoplasmic (S6K, 4E-BP1) substrates of mTORC1 upon BafA1 treatment (1-8 h). Note the rapid drop in TFEB phosphorylation, whereas that of S6K/4E-BP1 remains largely unaffected even at much later time points (c). Quantification of TFEB phosphorylation in (d). Quantification of S6K phosphorylation in (e). n = 3 independent experiments. **(f-g)** The rate of S6K dephosphorylation is similar between Rag-proficient and Rag-deficient cells. Rapamycin time course (1-30 min) in control (WT) and RagA/B knockout (KO) cells, assessing S6K dephosphorylation kinetics (f). Quantification of S6K phosphorylation in (g). n = 3 independent experiments. Arrowheads indicate bands corresponding to different protein forms, when multiple bands are present. P: phosphorylated form. Data in graphs shown as mean ± SEM. ** p<0.01, *** p<0.001, **** p<0.0001, ns: non-significant.

In sum, using an orthogonal validation scheme, our data from five independent pharmacological and genetic perturbations that block basal lysosomal protein degradation i) indicate the local AA production as the reason why a proportion of mTOR localizes to lysosomes, ii) reveal that non-lysosomal mTOR retains its activity toward its canonical substrates, but not toward its lysosomal substrates, under basal culture conditions, and iii) show that the activity of this non-lysosomal mTORC1 pool is still regulated by exogenous AA availability.

### The lysosomal Rag GTPases are substrate- and location-specific regulators of mTORC1

Because mTORC1 recruitment to lysosomes is mediated by the Rag GTPases, to genetically dissect the lysosomal from the non-lysosomal mTORC1 regulation, and to study the functional consequences of lysosomal delocalization of mTORC1, we generated various Rag loss-of-function cell line models, lacking RagA/B or RagC/D expression (Fig. 5a). As expected ^19,23,24^, the lysosomal mTOR localization was diminished in RagA/B KO cells, under all nutritional conditions (Fig. 5b,c). To independently validate our microscopy data, we generated control or RagA/B KO cells stably expressing HA-tagged TMEM192 as a lysosomal anchor (or FLAG-tagged TMEM192 as a negative control), and developed a modified anti-HA lyso-IP protocol (original version described in ^31^) that allowed us to isolate fractions enriched for intact lysosomes as well as non-lysosomal fractions. These experiments confirmed that a fraction of mTORC1 (shown by the presence of mTOR and Raptor) is specifically co-purified with lysosomes in a Rag-dependent manner, while both wild-type (WT) and RagA/B KO cells contain non-lysosomal mTORC1 complexes (Fig. 5d).

**Figure 5.**
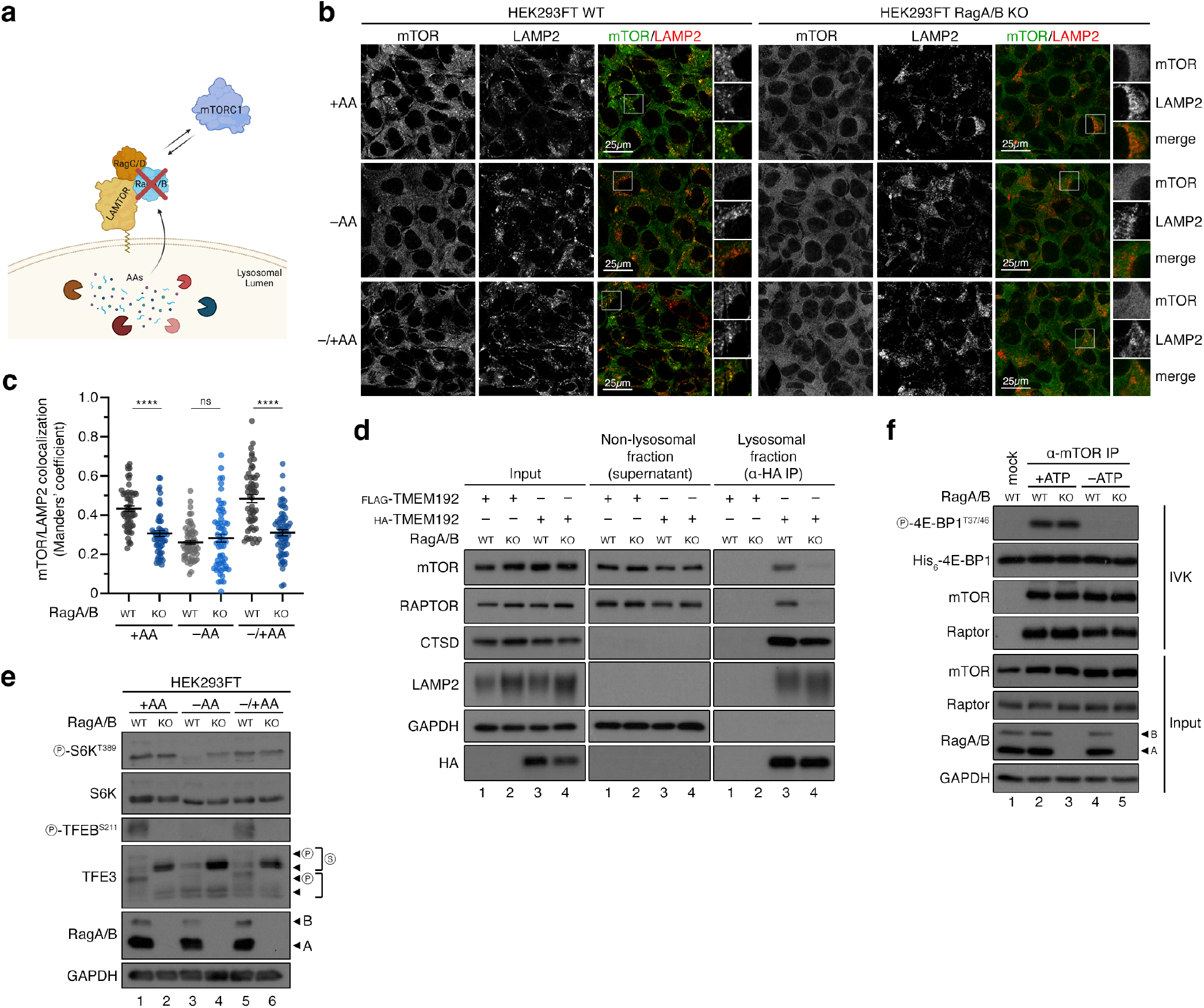
Rag knockout cells have non-lysosomal but active mTORC1 toward canonical substrates. **(a)** Schematic model for the genetic removal of the Rag GTPases. **(b-c)** Colocalization analysis of mTOR with LAMP2 (lysosomal marker) in HEK293FT WT and RagA/B KO cells, treated as indicated in the figure, using confocal microscopy. Scale bars = 25 μm. Magnified insets shown to the right (b). Quantification of colocalization in (c). n = 55-60 individual cells from 3-4 independent fields per condition. **(d)** Lyso-IP experiments with WT and RagA/B KO HEK293FT cells stably expressing HA-tagged TMEM192 (or FLAG-TMEM192 as negative control). Intact lysosomes were immunopurified by anti-HA IPs under native conditions, and the presence of LAMP2, cathepsin D (CTSD), mTOR and Raptor proteins in the lysosomal and non-lysosomal fractions, as well as in whole cell lysates, was analyzed by immunoblotting as indicated. **(e)** Immunoblots with lysates from HEK293FT WT and RagA/B KO cells, treated with media containing or lacking AAs, in basal (+AA), starvation (–AA) or add-back (–/+AA) conditions, probed with the indicated antibodies. **(f)** Summary table of mTORC1 substrate properties and dependencies. **(g)** IVKs with mTORC1 immunopurified from WT and RagA/B KO HEK293FT cells and recombinant 4E-BP1 protein used as substrate. 4E-BP1 phosphorylation detected by immunoblotting. Arrowheads indicate bands corresponding to different protein forms, when multiple bands are present. P: phosphorylated form. Data shown are representative of 3 replicate experiments. Data in (c) shown as mean ± SEM. **** p<0.0001, ns: non-significant. See also Ext. Data Figures 3-7.

Interestingly, despite mTORC1 being non-lysosomal in RagA/B KO cells, phosphorylation of its canonical substrate S6K was largely unaffected in unchallenged cells, grown under basal conditions (Fig. 5e), similar to our results using cells with perturbed lysosomal function (Fig. 1-4 and Ext. Data Fig. 1-2). Confirming the well-established role of the Rags in the acute re-activation of mTORC1 toward S6K phosphorylation ^19,23^, RagA/B KO cells showed blunted recovery of mTORC1 activity upon AA re-supplementation, following AA depletion (Fig. 5e). Moreover, these cells demonstrated a partial resistance to AA starvation (Fig. 5e), which we and others have described previously ^24,32^, and is due to loss of the Rag-mediated lysosomal recruitment of TSC (Tuberous Sclerosis Complex; a negative regulator of mTORC1 activity) upon starvation ^24^. In contrast to the behaviour of canonical substrate phosphorylation, that of the lysosomal TFEB/TFE3 substrates was completely abolished in RagA/B KO cells under all conditions of AA availability (Fig. 5e). Further strengthening the Rag-dependent separation of mTORC1 activities, and showing that this is not a clonal artefact or a cell-type-specific characteristic of the RagA/B KO HEK293FT cells, similar data were obtained using RagC/D KO HEK293FT cells (Ext. Data Fig. 3a-c), RagA/B KO MEFs (Ext. Data Fig. 3d-f), and RagA/B KO SW-620 colorectal cancer cells (Ext. Data Fig. 3g). Moreover, the observed Rag- and AA-dependent effects were specific for mTORC1, as phosphorylation of AKT, a *bona fide* mTORC2 substrate was unaffected upon AA starvation or re-addition in control, RagA/B or RagC/D KO cells (Ext. Data Fig. 3h,i). Finally, to assess the intrinsic activity of mTORC1, we immunopurified endogenous mTOR complexes from control or RagA/B KO cells and performed *in vitro* kinase (IVK) assays, using recombinant 4E-BP1 as a substrate. Strikingly, not only the phosphorylation of the non-lysosomal mTORC1 substrates in cells, but also *in vitro* mTORC1 kinase activity was largely unaffected by Rag loss-of-function (Fig. 5f).

In support of our model that the Rags are involved primarily in the regulation of the lysosomal mTORC1 substrates, exogenous expression of an active-locked RagA (Q66L) mutant in RagA/B KO cells caused a striking increase in TFEB phosphorylation, while it only marginally increased the phosphorylation of S6K. Re-expression of WT RagA also rescued TFEB phosphorylation, albeit to a much lesser extent (Ext. Data Fig. 3j). These data are in line with a recent study from our group showing that stable reconstitution of a different Rag KO model with cancer-associated, activating RagC mutants (as a dimer with RagA) strongly increased TFEB/TFE3 phosphorylation, without affecting that of S6K (Fig. 5 in ^22^). In sum, neither loss nor re-expression of the Rags influences mTORC1 activity toward S6K, while TFEB phosphorylation is fully dependent on the presence and the activation status of the Rag dimer.

The Rag GTPase dimer is tethered to the lysosomal surface indirectly, via protein-protein interactions with the LAMTOR complex (also referred to as ‘Ragulator’) ^23,33^. Therefore, we next transiently knocked down the LAMTOR1/p14 subunit of the LAMTOR complex (Ext. Data Fig. 4a) as an additional means to dissociate mTORC1 from lysosomes, without perturbing the Rag dimer itself (Ext. Data Fig. 4b). As expected, siLAMTOR1 cells showed diffuse cytoplasmic localization of mTOR with no lysosomal accumulations (Ext. Data Fig. 4c,d). In agreement with our findings from Rag KO cells, *LAMTOR1* knockdown strongly diminished the phosphorylation of TFEB, without affecting S6K or 4E-BP1 phosphorylation under basal culture conditions (Ext. Data Fig. 4e).

In contrast to TFEB/TFE3, the canonical mTORC1 substrates such as S6K and 4E-BP1 were previously shown not to localize to lysosomes ^27,34^ and are generally considered to be cytoplasmic mTORC1 substrates (with S6K partly also localizing in the nucleus ^35^). Indeed, by lyso-IP experiments, we confirmed that S6K is found in the non-lysosomal fraction and does not co-purify with lysosomes of WT or RagA/B KO cells (Fig. 6a), in contrast to phosphorylated TFEB which is found strongly enriched in the lysosomal fraction specifically of WT cells, but not Rag KOs. The absence of S6K localization to lysosomes is also supported by a previous proteomic study showing that the vast majority of p70-S6K1 interaction partners are cytosolic (or nuclear) proteins, and not associated with lysosomes ^36^. In sum, our data from multiple independent approaches show that both mTOR and S6K localize to the cytoplasm of Rag-deficient cells, where S6K phosphorylation likely takes place. This is consistent with the role of S6K in protein synthesis, a process that takes place primarily in the cytoplasm.

**Figure 6.**
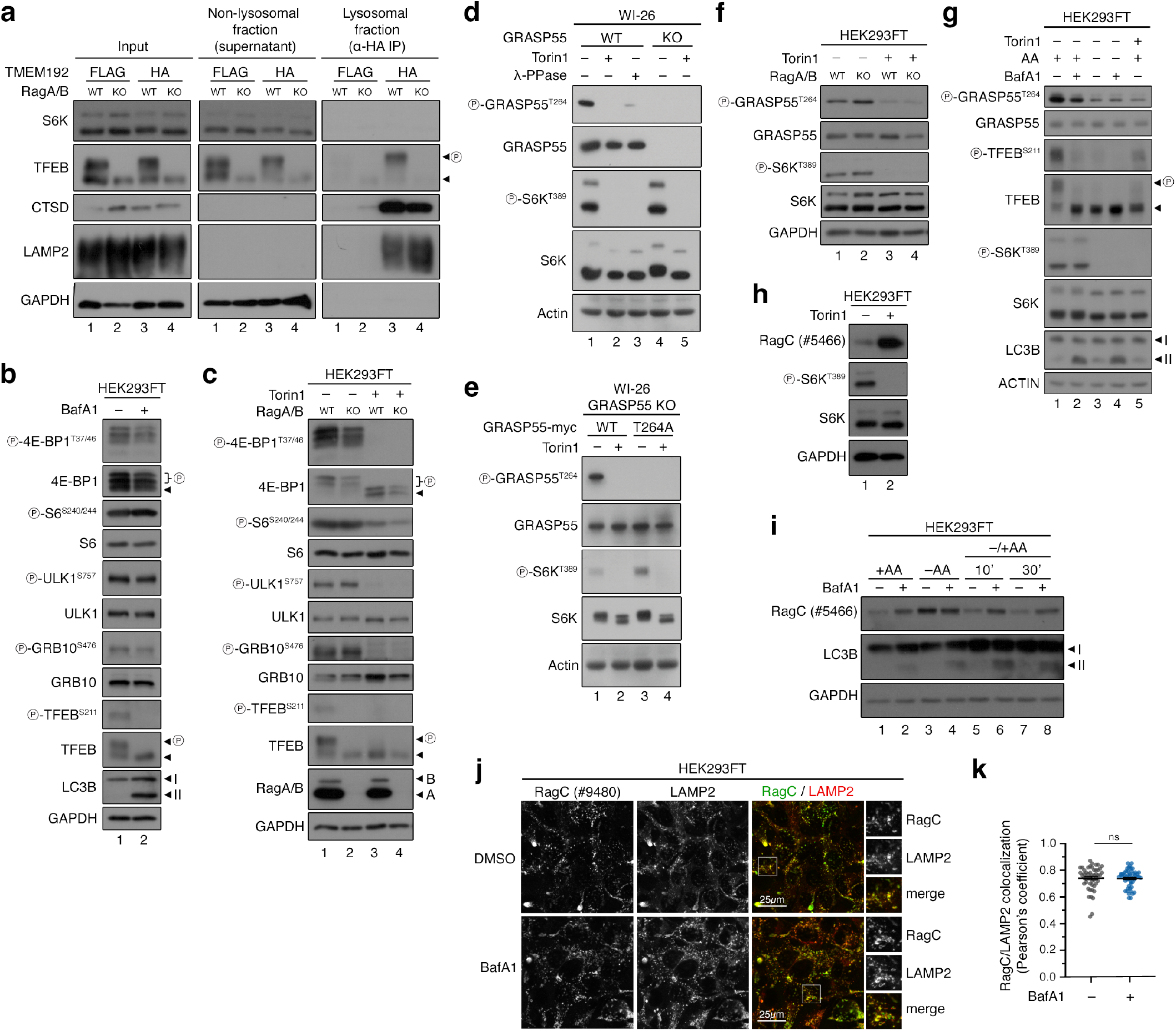
Characterization of mTORC1 activity toward additional lysosomal and non-lysosomal substrates. **(a)** Lyso-IP experiments in WT and RagA/B KO HEK293FT cells stably expressing HA-tagged TMEM192 (or FLAG-TMEM192 as negative control). Intact lysosomes were immunopurified by anti-HA IPs under native conditions, and the presence of LAMP2, cathepsin D (CTSD), TFEB and S6K proteins in the lysosomal and non-lysosomal fractions, as well as in whole cell lysates, was analyzed by immunoblotting as indicated. Note the absence of S6K from lysosomal fractions, and the presence of phospho-TFEB in the lysosomal fractions only of control cells. n = 2 independent experiments. (**b-c**) The phosphorylation of multiple mTORC1 substrates is largely unaffected by BafA1 treatment (b) or loss of Rag GTPases (c). In (c), Torin1 was used as a control for mTOR inhibition. n = 3 independent experiments. **(d-e)** Characterization of the p-GRASP55 antibody. The signal from p-GRASP55 on T264 is diminished by Torin1 or λ-phosphatase (λ-PPase) treatment, and is lost in WI-26 cells not expressing GRASP55 (d). GRASP55 KO WI-26 cells reconstituted with WT or T264A mutant GRASP55 (myc-tagged), and treated with Torin1, or DMSO as control. Note that p-GRASP55 signal is only detected in cells expressing WT GRASP55 with active mTORC1 (e). **(f-g)** GRASP55 phosphorylation by mTORC1 is retained in RagA/B KO (f) or BafA1-treated cells (g), similarly to that of S6K. n = 3 independent experiments. **(h)** Downregulation of RagC phosphorylation by treatment with Torin1, shown as an increase in RagC signal detected with the #5466 anti-RagC antibody. n = 2 independent experiments. **(i)** RagC is an additional lysosomal mTORC1 substrate that requires properly-functioning lysosomes for its phosphorylation, similarly to TFEB/TFE3. Amino acid (AA) starvation or blockage of lysosomal function with BafA1 decrease RagC phosphorylation (shown as elevated RagC signal with #5466). n = 3 independent experiments. **(j-k)** Lysosomal localization of RagC is unaffected by BafA1 treatment (j). Quantification of RagC/LAMP2 colocalization in (k). n = 50 individual cells from 5 independent fields per condition. Representative data from one out of two independent experiments are shown. Arrowheads indicate bands corresponding to different protein forms, when multiple bands are present. P: phosphorylated form. Data in (k) shown as mean ± SEM. ns: non-significant.

In line with our data on S6K and 4E-BP1, the phosphorylation of additional non-lysosomal mTORC1 substrates and downstream effectors like ULK1, GRB10, and S6 was largely unaffected in BafA1-treated (Fig. 6b) or RagA/B KO cells (Fig. 6c), two conditions where mTOR is non-lysosomal and the lysosomal substrates like TFEB are completely dephosphorylated. GRASP55 (Golgi re-assembly and stacking protein 55) is a Golgi-residing protein that functions as a molecular switch for unconventional protein secretion (UPS), a cellular process activated upon starvation or stress. We have recently demonstrated that, when active, mTORC1 directly phosphorylates GRASP55 on Thr264 at the Golgi to inhibit UPS ^37^. As with the other non-lysosomal substrates, the specific GRASP55 phosphorylation by mTORC1 was not affected by loss of the Rags, and only mildly affected by BafA1 treatment, as assayed with a custom-made phospho-specific antibody (Fig. 6d-g). Finally, a recent study identified RagC as a direct mTORC1 substrate ^38^. As its primary localization is on the lysosomal surface, we tested whether RagC phosphorylation requires lysosomal function or it behaves similarly to the cytoplasmic substrates. To achieve this, we made use of an anti-RagC antibody that is raised against the N-terminus of the protein and preferentially recognizes non-phosphorylated RagC, therefore the signal in immunoblots anti-correlates with RagC phosphorylation. Accordingly, either Torin1 treatment (Fig. 6h) or AA starvation (Fig. 6i, compare lanes 1 and 3) increased RagC signal. Notably, blocking lysosomal function and inhibiting lysosomal mTORC1 activity with BafA1 also led to a strong increase in RagC signal, indicating strongly decreased RagC phosphorylation (Fig. 6i), similarly to what we observed for the other lysosomal mTORC1 substrates TFEB and TFE3. The dephosphorylation of RagC occurred despite the fact that its localization to lysosomes remained unaffected by BafA1 treatment (Fig. 6j,k), which indicates this is due to the delocalization of the kinase (i.e., mTORC1) and not the substrate (i.e., RagC) away from the lysosomal surface.

Overall, these data showed that, not only the Rags, but the complete lysosomal mTORC1 recruitment machinery is important for the regulation of the lysosomal mTORC1 substrates, and that the spatial and functional separation of mTORC1 activities can be achieved, not only by perturbing the expression of the Rags, but also their anchoring to the lysosomal membrane. Moreover, we find that the Rags are largely dispensable for basal mTORC1 activity in unchallenged cells, while the re-activation of mTORC1 is Rag-dependent, as reported previously ^19,24^. Finally, the Rag GTPases moonlight as substrate-specific regulators of mTORC1.

### Non-lysosomal mTORC1 activation by specific exogenous AAs occurs independently from known Rag-related signalling network components

Using multiple genetic and pharmacological ways to target the lysosomal AA sensing machinery, we dissociated the lysosomal localization of mTORC1 from its activity, revealing that non-lysosomal mTORC1 remains active toward its non-lysosomal substrates, including S6K and 4E-BP1. Indeed, by treating RagA/B or RagC/D KO cells with Torin1, a catalytic mTOR inhibitor, we confirmed that this persistent S6K phosphorylation is indeed mTOR kinase-activity-dependent (Ext. Data Fig. 5a,b). As mentioned above, albeit partially resistant at the early time points of AA starvation ^24^, these cells do respond to AA starvation, with complete mTORC1 inactivation achieved at slightly later time points, thus showing that non-lysosomal mTORC1 can be regulated by exogenous AA availability (Ext. Data Fig. 5a-c).

The TSC/Rheb signalling hub lies directly upstream of mTORC1, integrating information from most stimuli that regulate mTORC1 activity, including AAs and growth factors ^3,24,39–41^. Transient downregulation of TSC2 in control or RagA/B KO cells—that leads to hyperactivation of Rheb—robustly elevated mTORC1 activity in both genotypes (Ext. Data Fig. 5d), showing that these upstream regulators are relevant also for the activity of non-lysosomal mTORC1 toward S6K. Consistent with this, blocking growth factor signalling by using a pharmacological Akt inhibitor decreased S6K phosphorylation in RagA/B KO and control cells to a similar extent (Ext. Data Fig. 5e). Accordingly, serum starvation and FBS or insulin re-supplementation experiments in Rag KO MEFs confirmed that mTORC1 activity properly responds to growth factor or insulin stimulation in Rag-deficient cells (Ext. Data Fig. 5f), as in the respective Rag-proficient cells (Ext. Data Fig. 5g). However, in line with previous work on the role of the Rags in glucose signalling to mTORC1 on lysosomes (see e.g., Fig. 2C in ^32^), its activity (assayed by the phosphorylation of S6K and 4E-BP1) did not respond to glucose starvation (Ext. Data Fig. 5h, compare lanes 6 & 7), while it was readily downregulated by AA starvation (Ext. Data Fig. 5h, compare lanes 6 & 9). Importantly, this demonstrates that non-lysosomal mTORC1 specifically senses the absence of exogenous AAs, but not of glucose.

The fact that the non-lysosomal activity of mTORC1 is under the control of exogenous AA sources hints at the existence of Rag- and lysosome-independent regulatory mechanisms via which different AA pools signal to mTORC1. The Sestrin1/2 and CASTOR1/2 proteins are cytoplasmic sensors for leucine and arginine, respectively, signalling AA availability to the Rag GTPases via the GATOR2 complex ^42–47^ (Ext. Data Fig. 6a). Therefore, we performed transient knockdown experiments targeting Mios, a key component of the GATOR2 protein complex, as a tool to study the possible involvement of the cytoplasmic AA sensors in the regulation of non-lysosomal mTORC1. Although downregulation of Mios strongly decreased lysosomal mTOR localization (Ext. Data Fig. 6b,c), consistent with its previously described role in AA sensing upstream of the Rags ^48,49^, and suppressed TFEB phosphorylation under all nutritional conditions, the phosphorylation of S6K and 4E-BP1 was unaffected in cells grown under basal conditions (Ext Data Fig. 6d). As observed also for cells with Rag loss-of-function, Mios knockdown potently blunted the re-activation of mTORC1 upon AA re-addition (Ext Data Fig. 6d). Therefore, although this complex was described to integrate information from proteins that sense AA sufficiency in the cytoplasm, it still signals through the lysosomal Rag-related machinery to regulate mTORC1. These data suggest that extracellular AAs likely signal to non-lysosomal mTORC1 via mechanisms that have not been resolved yet. Consistent with this hypothesis, we found that RagA/B KO cells respond to different AA groups, compared to their wild-type counterparts. In particular, while mTORC1 activity in Rag-proficient control cells is sensitive to depletion of hydrophobic (methionine, leucine, isoleucine, glycine, valine; ‘– MLIGV’) or positively-charged (histidine, arginine, lysine; ‘–HRK’) AAs from the culture media (Fig. 7a,b), RagA/B KO cells do not respond significantly to removal of these AA groups (Fig. 7c,d). Interestingly, treatment with starvation media specifically lacking serine, threonine and cysteine (‘–STC’) downregulated mTORC1 activity in RagA/B KO, but not in control cells (Ext. Data Fig. 7a-d). In addition, each of the AAs within the ‘–STC’ mix was necessary for mTORC1 to stay active in Rag KO cells, as treating cells with media lacking serine, threonine, and—to a lesser extent—cysteine, singly, downregulated mTORC1 comparably to the combined removal of S+T+C (Fig. 7e,f). Because Rag-deficient cells (in which mTORC1 is non-lysosomal) demonstrate differential sensitivity of mTORC1 activity to distinct AA subgroups—compared to Rag-proficient cells that show predominantly lysosomal mTORC1— it is intriguing to predict that different sensing mechanisms exist to regulate mTORC1 in the two subcellular locations. Importantly, whereas the lysosomal Rag-related AA sensing machinery has been described extensively, the proteins and pathways that signal the availability of specific exogenous AAs to non-lysosomal mTORC1 are completely unknown. Future endeavours will be necessary to put together this AA sensing network.

**Figure 7.**
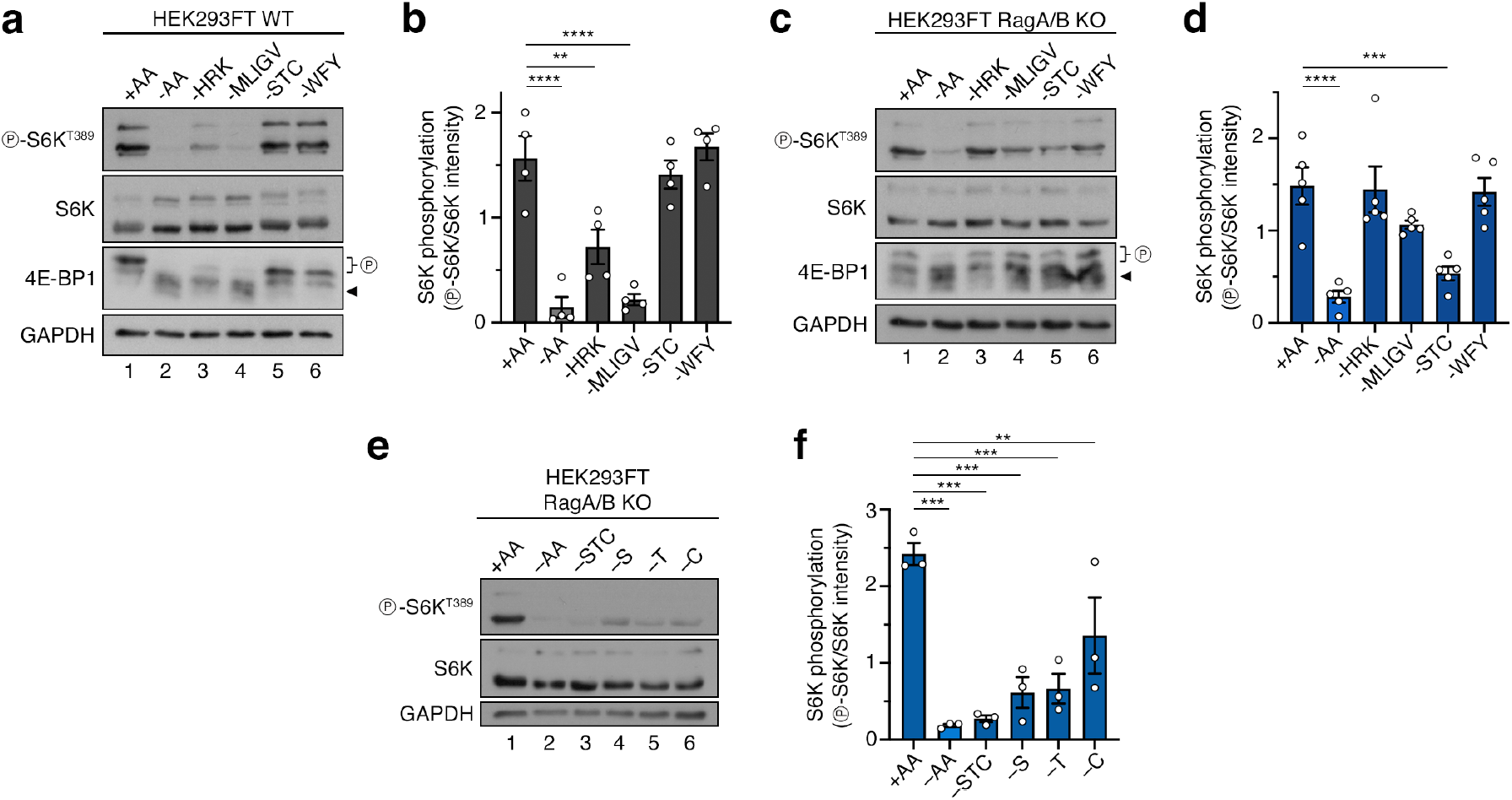
Non-lysosomal mTORC1 is regulated by distinct exogenous AAs. **(a-b)** Immunoblots with lysates from HEK293FT WT cells, treated with media containing or lacking the AA groups shown in the panel, probed with the indicated antibodies (a). Quantification of mTORC1 activity (p-S6K^T389^/S6K) in (b). n = 4 independent experiments. **(c-d)** As in (a-b), but for RagA/B KO HEK293FT cells (c). Quantification of mTORC1 activity (p-S6K^T389^/S6K) in (d). n = 5 independent experiments. **(e-f)** As in (c-d), but for RagA/B KO HEK293FTs treated with media lacking the indicated groups or individual AAs (e). Quantification of mTORC1 activity (p-S6K^T389^/S6K) in (f). n = 3 independent experiments. Arrowheads indicate bands corresponding to different protein forms, when multiple bands are present. P: phosphorylated form. Data in graphs shown as mean ± SEM. ** p<0.01, *** p<0.001, **** p<0.0001.

Glutamine and asparagine re-supplementation was recently reported to re-activate mTORC1 (toward its canonical substrates) via the Golgi-localized Arf1 GTPase and independently from the Rags ^50,51^ (Ext. Data Fig. 7a). Therefore, we next tested if Arf1 is involved in the regulation of non-lysosomal mTORC1, by performing transient knockdown experiments in RagA/B KO cells (Ext. Data Fig. 7b). Although Arf1 was indeed important for full re-activation of mTORC1 upon AA add-back, it did not influence its activity under basal conditions (Ext. Data Fig. 7c). Similarly, treatment with Golgicide A (GA) or Brefeldin A (BFA), two drugs that target the ArfGEF GBF1 (Ext. Data Fig. 7a), blocked Arf1 activation, as shown by the robust structural and morphological changes on the Golgi apparatus (Ext. Data Fig. 7d). These treatments did not alter mTORC1 activity in either RagA/B KO or control cells, toward any of its substrates (Ext. Data Fig. 7e). Therefore, like the Rags that are dispensable for basal mTORC1 activity, Arf1 is seemingly also involved only in the re-activation of mTORC1 upon re-supplementation with specific AAs, like e.g., glutamine. Similar results were obtained previously in yeast cells, in which glutamine can regulate TORC1 in the absence of the yeast Rag homologs ^52^.

### Lysosomal and non-lysosomal mTORC1 complexes regulate different cellular processes via the phosphorylation of distinct effectors

We describe here the spatial separation of mTORC1 activities toward different substrates and downstream effectors. Therefore, we next sought to investigate what is the physiological function of these two distinct mTORC1 entities in cells. Because mTORC1 is known to regulate protein synthesis via the phosphorylation of S6K and 4E-BP1, we first assessed *de novo* protein synthesis by using a modified puromycin incorporation assay (OPP assay), comparing RagA/B KO to control cells. Consistently with S6K and 4E-BP1 phosphorylation being largely unaffected in *Rag*-null cells grown in AA-replete media, *de novo* protein synthesis was independent of Rag presence (Fig. 8a,b and Ext. Data Fig. 8a,b). Therefore, mTORC1 regulates protein synthesis independently of the Rag GTPases and the lysosomal amino acid sensing machinery.

**Figure 8.**
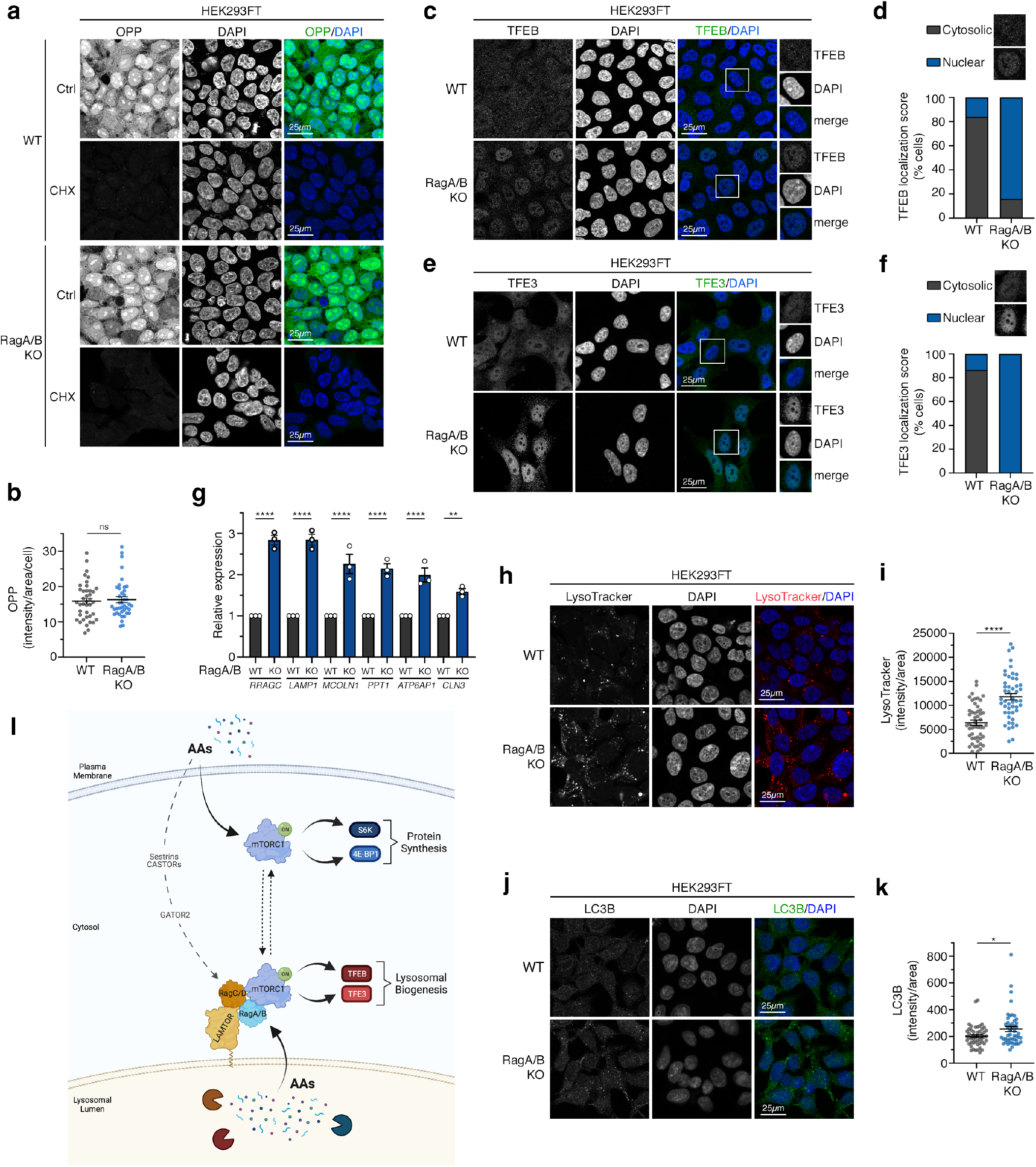
Functional separation of Rag-dependent and -independent mTORC1 activities. **(a-b)** *De novo* protein synthesis (OPP incorporation) assay with WT and RagA/B KO HEK293FT cells. Nuclei stained with DAPI. Cycloheximide (CHX) inhibitor used as negative control. Scale bars = 25 μm (a). Quantification of OPP signal in (b). n = 50 individual cells from 5 independent fields per condition. **(c-d)** TFEB localization analysis in WT and RagA/B KO HEK293FT cells, using confocal microscopy. Nuclei stained with DAPI. Magnified insets shown to the right. Scale bars = 25 μm (c). Scoring of TFEB localization in (d). Individual cells were scored for nuclear or cytoplasmic TFEB localization as indicated in the example images. n_WT_ = 65 cells, n_ABKO_ = 102 cells. **(e-f)** As in (c-d) but for TFE3 localization. Scoring (n_WT_ = 52 cells, n_ABKO_ = 52 cells) in (f). **(g)** Expression analysis of TFEB/TFE3 target genes in HEK293FT WT and RagA/B KO cells. Data shown as mean ± SD. **(h-i)** LysoTracker staining in HEK293FT WT and RagA/B KO cells. Nuclei stained with DAPI. Scale bars = 25 μm (h). Quantification of LysoTracker signal in (i). n = 50 individual cells from 5 independent fields per condition. **(j-k)** LC3B staining in HEK293FT WT and RagA/B KO cells. Nuclei stained with DAPI. Scale bars = 25 μm (j). Quantification of LC3B signal in (k). n = 50 individual cells from 5 independent fields per condition. **(l)** Working model for the functional separation of mTORC1 activities in cells. See main text for details. Created with BioRender.com. Data shown are representative of 3 replicate experiments. Data in (b), (g), (i), (k) shown as mean ± SEM. * p<0.05, ** p<0.01, **** p<0.0001, ns: non-significant. See also Ext. Data Figure 8.

Unlike for S6K and 4E-BP1, the mTORC1-dependent phosphorylation of the TFEB and TFE3 transcription factors is abolished in Rag-deficient cells. Accordingly, dephosphorylation of TFEB and TFE3 in RagA/B KO cells was accompanied by increased nuclear localization of these transcription factors (Fig. 8c-f). Similar data were obtained for TFE3 localization in RagC/D KO cells (Ext. Data Fig. 8c,d). In the nucleus, TFEB/TFE3 upregulated the expression of target genes that are related to lysosome and autophagosome biogenesis (Fig. 8g), in agreement with previous studies ^53–56^. Indeed, RagA/B KO HEK293FT cells showed an overall increase in lysosome abundance (assessed by LysoTracker staining) (Fig. 8h,i) as well as increased autophagosome content (shown by LC3 immunofluorescence) (Fig. 8j,k). Similar results were obtained in MEFs, with RagA/B KO cells demonstrating elevated LC3 levels (Ext. Data Fig. 8e,f).

A vast gene expression program is regulated downstream of mTORC1, directly or indirectly, via the activity of transcription factors (TFs) like SREBP, HIF1 ^57^, and ATF4 ^58^. To investigate whether this mTORC1-dependent gene expression program depends on the presence of the Rags, we performed RNA-seq experiments and tested the expression of selected target genes for each of these TFs, comparing WT and RagA/B KO HEK293FT cells. As a control, we also re-analysed our recently published RNA-seq dataset from Rapamycin-vs. DMSO-treated HEK293FT cells, in which mTORC1 is inhibited and S6K is dephosphorylated ^59^. These analyses showed that selected SREBP, HIF1 and ATF4 target genes were consistently downregulated upon mTORC1 inhibition with rapamycin, confirming that the activity of these TFs is regulated downstream of mTORC1 (Ext. Data Fig. 8g-i). In contrast, the expression of these target genes was not consistently affected in Rag KO cells (Ext. Data Fig. 8g-i), indicating that these transcription factors do not require the activity of lysosomal mTORC1, and thus are likely regulated by non-lysosomal mTORC1 entities. As a control, we expanded our qPCR analyses described above (Fig. 8g) to check the expression of additional TFEB targets in the two RNA-seq datasets. As expected from the loss of TFEB phosphorylation and its nuclear translocation in Rag KO cells, also the expression of its target genes was strongly upregulated (Ext. Data Fig. 8j). Moreover, consistent with TFEB being a rapamycin-resistant substrate of mTORC1 ^60^, expression of these TFEB targets was largely unaffected in rapamycin-treated cells (Ext. Data Fig. 8j). These data highlight the mTORC1-dependent transcriptional program as an additional physiological cellular process that is specifically controlled by non-lysosomal (e.g., for SREBP, HIF1, and ATF4 targets) or lysosomal mTORC1 (e.g., for TFEB targets). Taken together, we report here that the lysosomal and non-lysosomal mTORC1 entities regulate distinct cellular functions via the differential phosphorylation of distinct mTORC1 substrates at these subcellular locations (Fig. 8l).

## Discussion

The reason for a lysosome-centered regulation of mTORC1 has been a major field of discussion since its introduction in 2008 (discussed in ^4^). mTORC1 re-activation ‘makes sense’ to happen at lysosomes, where bulk macro-autophagy takes place upon amino acid starvation, degrading proteins to supply cells with fresh AAs, which in turn re-activate mTORC1. Therefore, mTORC1 is meaningfully localized in the vicinity of AA re-supplementation—following nutrient starvation—and posed for re-activation. However, this model does not sufficiently explain why mTORC1 should be activated on lysosomes in non-starved, unchallenged cells. Indeed, although the Rag GTPases localize primarily on lysosomes and regulate mTORC1 by recruiting it there ^23^, recent studies by us and others provide evidence that the lysosomal localization of mTORC1 and its activity (assayed by phosphorylation of S6K in most cases) are not always interrelated ^22,24,27,50,61–64^. Consistent with this, its direct regulators (e.g., Rheb, Rags, Arf1) ^27,28,50,65–70^, many of its substrates ^27,71–73^, and mTOR itself ^27,28,37,67,69,72,74–78^ were also described to localize at multiple subcellular locations, besides lysosomes ^4^. Furthermore, the relocalization of mTORC1 to lysosomes in response to AA re-supplementation is seemingly transient, dynamic, and involves only a fraction of the total mTOR cellular pool ^27,28^. Therefore, our study supports a model whereby the regulation of mTORC1 signalling by AAs also involves non-lysosomal locations and, thus, Rag-independent mechanisms that function in parallel to the lysosome-based, Rag-related AA-sensing network (Fig. 8l). Based on these previous data, we hypothesize that these mTORC1 ‘pools’ are not strictly separated from each other in wild-type cells, but instead the complexes likely dynamically relocalize between the lysosomal surface, the cytoplasm and presumably other organelles (e.g., the Golgi) too, in response to local AA stimuli. Such a spatial separation of mTORC1 regulation and activities, with mTORC1 phosphorylating different substrates at distinct, functionally-relevant subcellular locations, would intuitively confer specificity and compartmentalization of its function.

Previous attempts to identify novel regulators of dTOR (*Drosophila* TOR) activity in a genome-wide manner in untreated *Drosophila* cells, using the phosphorylation of S6 (a direct S6K substrate) as a read-out, failed to identify any Rag GTPase or LAMTOR orthologs among the hits ^79^. Furthermore, animal models lacking Rag activity also support the notion that additional, Rag-independent mechanisms play an important role in regulating mTORC1. According to the previously described mouse models, RagA-null animals die around embryonic day E10.5 ^32^. Moreover, LAMTOR2-null mouse embryos, which lose proper Rag localization and activity, also die shortly before E10.5 ^80^. In contrast, mTOR and Raptor knockouts die much earlier in embryogenesis (ca. E3.5 ^81,82^ and E6.5 ^83^, respectively), indicating that Rag loss-of-function does not phenocopy mTORC1 inactivation. In line with this, MEFs extracted from RagA-null mice grow similarly to their WT counterparts and exhibit persistent mTORC1 activity (assayed by phosphorylation of its canonical substrates), despite having blunted mTORC1 lysosomal localization ^32^. In another mouse model, deletion of both RagA and RagB slightly reduced (approx. 30%) but did not abolish mTORC1 activity in MEFs ^50^ or in cardiomyocytes ^84^. Genetic studies in zebrafish also showed that TORC1 signalling is apparently normal in RagA-mutant larvae ^85^. Hence, mTORC1 is active also in the absence of RagA/B, and therefore the Rags seem largely dispensable for physiological mTORC1 activation.

Some of these previous studies interpreted the persistent phosphorylation of S6K in Rag-deficient cells or animals as the result of compensatory activation of other signalling pathways that lead to mTORC1 activation. Our findings, however, show that this is not the case, for several reasons: i) the sustained S6K phosphorylation is indeed dependent on mTOR activity (Ext. Data Fig. 5a,b); ii) transient LAMTOR knockdown mimics the effects of chronic Rag depletion (Ext. Data Fig. 4); iii) short-term pharmacological interventions that perturb lysosomal function and local AA production and efflux are also sufficient to recapitulate the Rag KO phenotype (Fig. 1,2 and Ext. Data Fig. 2); iv) the phosphorylation of the lysosomal mTORC1 substrates (i.e., TFEB, TFE3, RagC) is actually diminished in all pharmacological or genetic models that we use here, therefore mTORC1 is selectively active toward its non-lysosomal substrates; v) transient re-expression of active RagA is sufficient to rescue lysosomal activity of mTORC1 in RagA/B KO cells (Ext. Data Fig. 3j); and vi) mTORC1 activity still responds properly to exogenous AA availability (Fig. 1-3, 5-7 and Ext. Data Fig. 2-3, 5-7) and to growth factor signalling (Ext. Data Fig. 5d-g) in cells where these complexes are non-lysosomal. Therefore, we propose that the active pools of non-lysosomal mTORC1 are physiologically relevant in cells and likely also in animal tissues.

If Rag KO mice maintain active mTORC1, why are they embryonically lethal? As cells with Rag loss-of-function do not completely respond to AA removal ^24^ and show compromised mTORC1 re-activation upon AA add-back ^19,24^, one possibility is that this is due to improper response to AA starvation and re-supplementation. Alternatively, because *Rag*-mutant cells have constitutively activated TFEB/TFE3 signalling, and because these transcription factors play important roles in the metabolic and physiological response to starvation in mice ^86,87^, we speculate that the dysregulation of the TFEB/TFE3 branch is likely the most important reason why Rag removal leads to lethality in mice.

Previous work by the Zoncu lab indicated that depletion of the FLCN-FNIP2 complex (an upstream regulator of RagC) alters TFE3 localization, without affecting the phosphorylation of S6K and 4E-BP1 ^88^. Along the same lines, an independent study from the Ballabio group showed that the phosphorylation and localization of TFEB is specifically controlled by a FLCN-RagC/D complex in response to AAs, but does not require growth factor signalling to Rheb ^89^. Interestingly, a more recent publication from the Henske lab described the disconnection of mTORC1 substrate phosphorylation downstream of TSC: whereas S6K phosphorylation increases in cells lacking TSC, TFEB/TFE3 are dephosphorylated and translocate to the nucleus ^90^. Our findings, presented here, expand these previous observations and provide a mechanistic and cell-biological explanation for the substrate specificity downstream of mTORC1, which, we now show, is defined by the spatial separation of mTORC1 activities both at the lysosome and away from it. Our data are also in agreement with previous work in yeast by the De Virgilio group, suggesting that this phenomenon is evolutionarily conserved ^91^. Importantly, together with these studies, our findings underscore the need to redefine the term ‘mTORC1 activity’ in the first place: stating that mTORC1 is ‘active’ or ‘inactive’ is clearly not sufficient, and one would need to specify whether mTORC1 is active or inactive toward a particular target.

Because we find that: i) under basal conditions, the non-lysosomal pool of mTORC1 is regulated by exogenous AAs, while the lysosomal pool mostly responds to AAs coming from the lysosomal lumen; ii) the two mTORC1 pools respond to different AA groups; and iii) AAs signal to non-lysosomal mTORC1 independently of the lysosomal Rag-related machinery or the known cytoplasmic AA sensors, we propose that additional, currently unidentified proteins and pathways mediate AA signalling to mTORC1 away from lysosomes. As with the intricate, Rag-associated signalling network that has been put together over the last 15 years, we envisage that a similarly complex AA sensing machinery remains to be identified in the years to come, to complete the picture of how cell growth, metabolism and nutrient sensing are coordinated in cells.

## Methods

### Cell culture

All cell lines were grown at 37 °C, 5% CO_2_. Human female embryonic kidney HEK293FT cells (#R70007, Invitrogen; RRID: CVCL_6911), immortalized mouse embryonic fibroblasts (MEFs), and human male colorectal cancer SW-620 cells (#CCL-227, ATCC; RRID:CVCL_0547) were cultured in high-glucose Dulbecco’s Modified Eagle Medium (DMEM) (#41965-039, Gibco), supplemented with 10% fetal bovine serum (FBS) (#F7524, Sigma; #S1810, Biowest). Human male diploid lung WI-26 SV40 fibroblasts (WI-26 cells; CCL-95.1, ATCC; RRID, CVCL_2758) were cultured in DMEM/F12 GlutaMAX medium (#31331093, Gibco) containing 10% FBS. All media were supplemented with 1x penicillin– streptomycin (15140-122, Gibco).

HEK293FT cells were purchased from Invitrogen. Wild-type control and RagA/B KO immortalized MEFs were a kind gift of Kun-Liang Guan (described in ^50^). SW-620 cells were obtained from ATCC before the initiation of the project. The identity of the WI-26 cells was validated using the Short Tandem Repeat (STR) profiling service, provided by Multiplexion GmbH. The identity of the HEK293FT cells was validated by the Multiplex human Cell Line Authentication test (Multiplexion GmbH), which uses a single nucleotide polymorphism (SNP) typing approach, and was performed as described at www.multiplexion.de. No commonly misidentified cell lines were used in this study. All cell lines were regularly tested for *Mycoplasma* contamination, using a PCR-based approach and were confirmed to be *Mycoplasma*-free.

### Cell culture treatments

Amino acid (AA) starvation experiments were performed as described previously ^24,39^. In brief, custom-made starvation media were formulated according to the Gibco recipe for high-glucose DMEM, specifically omitting all or individual amino acids, or specific amino acid groups, as indicated in the figures. The media were filtered through a 0.22-μm filter device and tested for proper pH and osmolality before use. For the respective AA-replete (+AA) treatment media, commercially available high-glucose DMEM was used (#41965039, Thermo Fisher Scientific). All treatment media were supplemented with 10% dialyzed FBS (dFBS) and 1x Penicillin-Streptomycin (#15140-122, Gibco). For this purpose, FBS was dialyzed against 1x PBS through 3,500 MWCO dialysis tubing. For basal (+AA) conditions, the culture media were replaced with +AA treatment media 60-90 min before lysis or fixation. For amino-acid starvation (-AA), culture media were replaced with starvation media for 1 hour, unless otherwise indicated in the figure legends. For AA add-back experiments, cells were first starved as described above and then starvation media were replaced with +AA treatment media for 30 min, unless otherwise indicated in the figures.

For growth factor starvation experiments, complete culture media were replaced with FBS-free DMEM supplemented with 1x penicillin-streptomycin (#15140-122, Gibco) for 1 hour. For growth factor re-addition, cells were first starved for growth factors as described above and then 10% FBS (#F7524, Sigma; #S1810, Biowest) or 1 μM insulin (#I9278, Sigma) were added to the media for 30 minutes before harvesting. For glucose starvation experiments, cells were cultured for 1 hour in glucose-free DMEM (#11966025, Gibco) supplemented with 10% dFBS and 1x penicillin-streptomycin. For the respective control wells, the culture media were replaced with high-glucose DMEM containing 10% dFBS and 1x penicillin-streptomycin at the beginning of the experiment. For glucose re-addition samples, cells were first starved for glucose as described above and media were then replaced with glucose-containing media for another 30 min before harvesting.

For Bafilomycin A1 (#BML-CM110-0100, Enzo) treatments, the drug was added to a final concentration of 100 nM in the media for 6 hours before lysis or fixation, unless otherwise indicated in the figure legends. For concanamycin A (#C9705, Sigma) treatment, the drug was added to a final concentration of 100 nM in the media for 6 hours before lysis or fixation. Chloroquine (#C6628, Sigma) was added to the media to a final concentration of 50 μM for 6 hours before lysis or fixation. Treatment with E64 (#2935.1, Roth) and pepstatin A (#P5318, Sigma) to block lysosomal protease activity was performed by adding a combination of E64 (25 μM) and pepstatin A (50 μM) in the media for 16 hours before lysis or fixation. For experiments including treatments with +AA and –AA media, bafilomycin, concanamycin, chloroquine, or E64+PepA were kept also in the treatment media. For all experiments that did not include treatments with starvation media, the culture medium was refreshed 90 minutes before lysis, also including fresh inhibitors. To inhibit mTOR kinase activity, Torin1 (#14379, Cell Signaling Technology) was added in the culture media (final concentration 250 nM) for 1 hour (HEK293FT experiments) or 2 hours (WI-26 experiments). Specific mTORC1 inhibition was performed by adding rapamycin (#S1039; Selleckchem) directly to the culture media (final concentration 20nM) for the times indicated in the figure panels. Akt inhibition was achieved by addition of the Akt inhibitor VIII (#ENZ-CHM125, Enzo) in the culture media for 30 min (final concentration of 10 µM). Golgicide A (#345862, Sigma) and Brefeldin A (#BUF075, Biorad) were added in the culture media at final concentrations of 10 µM and 10 µg/ml, respectively, for 1 hour. For all drug treatments, DMSO (4720.1, Roth) was used as control.

### Antibodies

A list of all primary antibodies used in this study is found in Supplementary Table 1.

The H4B4 and ABL-93 antibodies against LAMP2, were obtained from the Developmental Studies Hybridoma Bank, created by the NICHD of the NIH and maintained at The University of Iowa, Department of Biology. H4B4 was deposited to the DSHB by August, J.T. / Hildreth, J.E.K. (DSHB Hybridoma Product H4B4) ^92^. ABL-93 was deposited to the DSHB by August, J.T. (DSHB Hybridoma Product ABL-93) ^93^.

### Plasmids and molecular cloning

The pETM-11-4E-BP1 vector, used to express His_6_-tagged 4E-BP1 in bacteria, was generated by PCR-amplifying human 4E-BP1 from cDNA (prepared from HEK293FT cells) using appropriate primers and cloned in the NcoI-NotI restriction sites of pETM-11. The respective pETM-11-RagC vector, used to produce bacterially-expressed His_6_-tagged RagC, was generated by PCR-amplifying human RagC from a pcDNA3-FLAG-RagC expression plasmid (described in ^24^) using appropriate primers and cloned in the NcoI-NotI restriction sites of pETM-11. For the GRASP55-myc expression vectors (pITR-TTP-GRASP55-myc-His), C-terminally myc- and His-tagged GRASP55 (wild-type and T264A mutant) was PCR-amplified from the respective pcDNA4/TO/hGRASP55-myc-His plasmids (described in ^37,94^) and cloned into the into the sleeping-beauty-based, doxycycline-inducible pITR-TTP vector ^95^ using the SfiI/NotI restriction sites. The pcDNA3-FLAG-hRagA WT and pcDNA3-FLAG-hRagA Q66L constructs were described previously ^24^. The pSpCas9(BB)-2A-Puro (PX459) V2.0 plasmid was purchased from Addgene (plasmid #62988; deposited by Feng Zhang) and described in ^96^. The pLJC6-3xHA-TMEM192 and pLJC6-2xFLAG-TMEM192 plasmids ^97^ were purchased from Addgene (plasmids #104434 and #104435; deposited by the Sabatini lab). All restriction enzymes were purchased from Fermentas/Thermo Scientific. The integrity of all constructs was verified by sequencing. All DNA oligonucleotides used in this study are listed in Supplementary Table 2.

### mRNA isolation, cDNA synthesis and quantitative real-time PCR

Total mRNA was isolated from cells using a standard TRIzol/chloroform-based method (#15596018, Thermo Fisher Scientific), according to manufacturer’s instructions. For cDNA synthesis, mRNA was transcribed to cDNA using the RevertAid H Minus Reverse Transcriptase kit (#EP0451, Thermo Fisher Scientific) according to manufacturer’s instructions. The cDNAs were diluted 1:10 in nuclease-free water and 4 µl of diluted cDNA were used per reaction, together with 5 µl 2x Maxima SYBR Green/ROX qPCR master mix (#K0223, Thermo Fisher Scientific) and 1 µl primer mix (2.5 µM of forward and reverse primers). Reactions were set in technical triplicates in a StepOnePlus Real-Time PCR system (Applied Biosystems). Relative gene expression was calculated with the 2^-ΔΔCt^ method, with *RPL13a* as an internal control, and normalized to the expression of the gene in the respective siCtrl or WT sample. All qPCR primers used in this study are listed in Supplementary Table 2.

### Plasmid DNA transfections

Plasmid DNA transfections in HEK293FT cells were performed using Effectene transfection reagent (#301425, QIAGEN), according to the manufacturer’s instructions. For the reconstitution of GRASP55 KO WI-26 cells, plasmid DNA transfections were performed using the X-tremeGENE HP DNA transfection reagent (#06366236001, Roche) in a 2:1 DNA/transfection reagent ratio according to the manufacturer’s protocol.

### Generation of stable cell lines

For the generation of stable cell lines expressing HA-tagged TMEM192 (lyso-IP lines) or FLAG-tagged TMEM192 (negative control lines for anti-HA lyso-IPs), WT HEK293FT cells were transfected using the respective expression vectors. Forty-eight hours post transfection, cells were selected with 3 μg/ml puromycin (#A11138-03, Thermo Fisher Scientific). Single-cell clones that express similar TMEM192 levels were used in lyso-IP experiments.

Stable cell lines expressing WT or T264A mutant GRASP55 were generated by reconstituting GRASP55 KO WI-26 cells using a doxycycline-inducible sleeping-beauty-based transposon system ^95^ as described previously ^37^. In brief, cells were transfected with the transposon-flanked pITR-TTP-GRASP55-myc-His plasmids (WT and T264A GRASP55) described above together with the transposase expressing pCMV-Trp vector. Twenty-four hours post-transfection, puromycin (2 μg/ml) was added to the medium and cells were selected for 5 days. Single-cell colonies were picked using cloning cylinders (#CLS31668, Sigma-Aldrich) and expanded. Clones that express similar levels of WT and T264 GRASP55 (leaky expression, in the absence of doxycycline) were used in follow-up experiments.

### Generation of knockout cell lines

The GRASP55 KO WI-26 cells were described previously ^37^. The HEK293FT RagA/B ΚΟ, RagC/D ΚΟ, GNPTAB KO, SW-620 RagA/B ΚΟ, HEK293FT HA-TMEM192 RagA/B KO and FLAG-TMEM192 RagA/B KO cell lines were generated using the pX459-based CRISPR/Cas9 method, as described elsewhere ^96^. The sgRNA expression vectors were generated by cloning appropriate DNA oligonucleotides (Supplementary Table 2) into the BbsI restriction sites of the pX459 vector (#62988, Addgene). An empty pX459 vector was used to generate matching control cell lines. In brief, transfected cells were selected with 3 μg/ml puromycin (#A11138-03, Thermo Fisher Scientific) 48 hours post transfection. Single-cell clones were generated by single cell dilution and knockout clones were validated by immunoblotting and functional assays.

### Gene silencing experiments

Transient knockdown of *GNPTAB, LAMTOR1, TSC2, MIOS* and *ARF1,* were performed using siGENOME (pool of 4) gene-specific siRNAs (Horizon Discoveries). An siRNA duplex targeting the *R. reniformis* luciferase gene (RLuc) (#P-002070-01-50, Horizon Discoveries) was used as control. Transfections were performed using 20 nM siRNA and the Lipofectamine RNAiMAX transfection reagent (#13778075, Thermo Fisher Scientific), according to the manufacturer’s instructions. Cells were harvested or fixed 72 hours post-transfection and knockdown efficiency was verified by immunoblotting or quantitative real-time PCR.

### Cell lysis and immunoblotting

For standard SDS-PAGE and immunoblotting experiments, cells from a well of a 12-well plate were treated as indicated in the figures, washed once with serum-free DMEM, and lysed in 250 μl of ice-cold Triton lysis buffer (50 mM Tris pH 7.5, 1% Triton X-100, 150 mM NaCl, 50 mM NaF, 2 mM Na-vanadate, 0.011 gr/ml beta-glycerophosphate), supplemented with 1x PhosSTOP phosphatase inhibitors (#04906837001, Roche) and 1x cOmplete protease inhibitors (#11836153001, Roche), for 10 minutes on ice. Samples were clarified by centrifugation (14000 rpm, 15 min, 4 °C) and supernatants transferred to a new tube. Protein concentration was determined using a Protein Assay Dye Reagent (#5000006, Bio-Rad). Normalized samples were boiled in 1x SDS sample buffer for 5 min at 95 °C (6x SDS sample buffer: 350 mM Tris-HCl pH 6.8, 30% glycerol, 600 mM DTT, 12.8% SDS, 0.12% bromophenol blue). For WI-26 lysates, cells from a well of a 6-well plate were treated as indicated in the figures and lysed in-well with 300 µl of ice-cold WI-26 lysis buffer (50 mM Tris-HCl pH 7.5, 0.5% Triton X-100, 150 mM NaCl, 0.1% SDS), supplemented with 1x PhosSTOP phosphatase inhibitors (#04906837001, Roche) and 1x cOmplete protease inhibitors (#11836153001, Roche). Samples were clarified by centrifugation (12000 x g, 15 min, 4 °C) and supernatants transferred to a new tube. Samples were boiled in 1x SDS sample buffer for 5 min at 95 °C.

For protein secretion experiments, WT or GNPTAB KO HEK293FT cells were cultured in serum-free media for 16 hours. Supernatants were collected and centrifuged (2000 x g, 5 min, 4 °C) to remove dead cells and debris. Cleared supernatants were concentrated using 3 kDa cut-off concentrator tubes (#516-0227P, VWR), according to the manufacturer’s instructions. SDS sample buffer (1x) was added to the concentrated supernatants and samples were boiled for 5 min at 95 °C before loading into SDS-PAGE gels.

Protein samples were subjected to electrophoretic separation on SDS-PAGE and analysed by standard Western blotting techniques. In brief, proteins were transferred to nitrocellulose membranes (#10600002 or #10600001, Amersham) and stained with 0.2% Ponceau solution (#33427-01, Serva) to confirm equal loading. Membranes were blocked with 5% skim milk powder (#42590, Serva) in PBS-T [1x PBS, 0.1% Tween-20 (#A1389, AppliChem)] for 1 hour at room temperature, washed three times for 10 min with PBS-T and incubated with primary antibodies [1:1000 in PBS-T, 5% bovine serum albumin (BSA; #10735086001, Roche)] rotating overnight at 4 °C. The next day, membranes were washed three times for 10 min with PBS-T and incubated with appropriate HRP-conjugated secondary antibodies (1:10000 in PBS-T, 5% milk) for 1 hour at room temperature. For the experiments using WI-26 samples or HEK293FT samples to assay GRASP55 phosphorylation upon BafA1 treatment, proteins were transferred to PVDF membranes (#10600023, Amersham). Equal loading was confirmed by staining with Ponceau S, blocked with 5% skim milk powder (#42590, Serva) in TBS-T buffer (50 mM Tris-HCl pH 7.4, 150 mM NaCl, 0.1% Tween-20) and incubated with primary antibodies diluted in TBS-T, for 1 h at room temperature or overnight at 4 °C, followed by incubation with appropriate HRP-conjugated secondary antibodies (1:10000 in TBS-T) for 1 h at room temperature. Signals were detected by enhanced chemiluminescence (ECL), using the ECL Western Blotting Substrate (#W1015, Promega); or SuperSignal West Pico PLUS (#34577, Thermo Scientific) and SuperSignal West Femto Substrate (#34095, Thermo Scientific) for weaker signals. Immunoblot images were captured on films (#28906835, GE Healthcare; #4741019289, Fujifilm). Blots were quantified using GelAnalyzer 19.1.

### Lambda-phosphatase treatment assays

Lambda-phosphatase (λ-PPase) treatment experiments were performed as one of the ways to validate the specificity of the custom-made phospho-specific antibody recognising GRASP55 phosphorylated on T264. In brief, cells were lysed in 300 μl ice-cold Triton lysis buffer (50 mM Tris-HCl pH 7.5, 0.5% Triton X-100, 150 mM NaCl, 0.1% SDS), supplemented with 1x EDTA-free cOmplete protease inhibitors (#11873580001, Roche), as described above. Lysates were cleared by centrifugation (15 min, 12000 x g) and 100 units of λ-phosphatase (#P0753, New England Biolabs) were added to the supernatants, followed by 30 min incubation at 30 °C. SDS sample buffer (1x final concentration) was added to the reactions, samples were boiled for 5 min at 95 °C and analysed by immunoblotting as described above.

### Lysosome purification (Lyso-IP) assays

To biochemically isolate intact lysosomes and associated proteins, we developed a modified lyso-IP method, based on the protocol previously described by the Sabatini group ^31^, which allowed us to also assess the non-lysosomal fractions. In brief, cells were seeded on a 15 cm dish until they reached 80-90% confluency, washed 2x with ice-cold PBS and scraped in 1 mL ice-cold PBS, containing 1x PhosSTOP phosphatase inhibitors (#04906837001, Roche) and 1x cOmplete protease inhibitors (#11697498001, Roche). Cells were then pelleted by centrifugation (1000 x g, 2 min, 4°C) and resuspended in 1 mL ice-cold PBS containing phosphatase and protease inhibitors. For input samples, 25 μl of the cell suspension were transferred in a new tube and lysed by the addition of 125 μl of Triton lysis buffer (50 mM Tris pH 7.5, 1% Triton X-100, 150 mM NaCl, 50 mM NaF, 2mM Na-vanadate, 0.011 gr/ml beta-glycerophosphate), supplemented with 1x PhosSTOP phosphatase inhibitors (#04906837001, Roche) and 1x cOmplete protease inhibitors (#11836153001, Roche) on ice for 10 min. Lysed input samples were then cleared by centrifugation (14000 x g, 15 min, 4°C) and the supernatant was transferred to new tubes containing 37.5 μl of 6x SDS sample buffer and boiled for 5 min at 95°C. For the lysosomal and non-lysosomal fractions, the remaining cell suspension was homogenized with 20 strokes in pre-chilled 2 mL hand dounce homogenizers kept on ice. The homogenate was cleared by centrifugation to remove unbroken cells (1000 x g, 2 min, 4°C) and the supernatant was incubated with 100 μl pre-washed Pierce anti-HA magnetic beads (#88837, Thermo Fisher Scientific) on a nutating mixer for 3 min at room temperature. After incubation with the beads, the supernatant was transferred to a new tube and centrifuged at high speed (20000 x g, 10 min, 4°C) to remove membranes and other organelles and retrieve the non-lysosomal/cytoplasmic fraction. Twenty-five microliters of the cleared supernatant were transferred in a new tube, mixed with 125 µl Triton lysis buffer, and incubated for 10 min on ice. Next, 37.5 µl 6x SDS sample buffer was added and samples were boiled. For the lysosomal fraction, beads were washed three times with ice-cold PBS containing phosphatase and protease inhibitors using a DynaMag spin magnet (#12320D, Invitrogen). After the last wash, lysosomes were eluted from the beads by addition of 50 µl Triton lysis buffer and incubation for 10 min of ice. Isolated lysosomes were then transferred to a new tube, 12.5 µl 6x SDS sample buffer was added and samples were boiled.

### Production of recombinant His_6_-tagged 4E-BP1 protein in bacteria

Recombinant His_6_-tagged 4E-BP1 protein was produced by transforming *E. coli* BL21 RP electrocompetent bacteria with the pETM-11-4E-BP1 vector described above, according to standard procedures. In brief, protein expression was induced with IPTG (isopropyl-β-D-thiogalactopyranoside) for 3 h at 30 °C, and His_6_-4E-BP1 was purified using Ni-NTA agarose (#1018244, QIAGEN) and eluted with 250 mM imidazole (#A1073, Applichem).

### mTORC1 kinase activity assays

*In vitro* mTORC1 kinase assays were developed based on previous reports ^98,99^, using endogenous mTORC1 complexes immunopurified from HEK293FT WT or RagA/B KO cells.

In brief, cells of a near-confluent 10 cm dish were lysed in CHAPS IP buffer (50 mM Tris pH 7.5, 0.3% CHAPS, 150 mM NaCl, 50 mM NaF, 2 mM Na-vanadate, 0.011 gr/ml beta-glycerophosphate), supplemented with 1x PhosSTOP phosphatase inhibitors (#04906837001, Roche) and 1x cOmplete protease inhibitors (#11836153001, Roche) for 10 min on ice. Samples were clarified by centrifugation (14000 rpm, 15 min, 4 °C), supernatants were collected and a portion was kept as input material. The remaining supernatants were subjected to immunoprecipitation by incubation with 2 μl of anti-mTOR antibody (#2983, Cell Signaling Technology) for 3 hours (4 °C, rotating), followed by incubation with 30 μl of pre-washed Protein A agarose bead slurry (#11134515001, Roche) for an additional hour (4 °C, rotating). Beads were then washed four times with CHAPS IP wash buffer (50 mM Tris pH 7.5, 0.3% CHAPS, 150 mM NaCl, 50 mM NaF) and once with kinase wash buffer (25 mM HEPES pH 7.4, 20 mM KCl), and excess liquid was removed with a Hamilton syringe. Kinase reactions were prepared by adding 10 μl 3x kinase assay buffer (75 mM HEPES/KOH pH 7.4, 60 mM KCl, 30 mM MgCl_2_) to the beads. Reactions were started by adding 10 μl of kinase assay start buffer (25 mM HEPES/KOH pH 7.4, 140 mM KCl, 10 mM MgCl_2_), supplemented with 500 μM ATP and 35 ng recombinant His_6_-4E-BP1 substrate. Reactions lacking ATP were set up as negative controls. All reactions were incubated at 30 °C for 30 min, and stopped by the addition of one volume 2x SDS sample buffer and boiling for 5 min at 95 °C. Samples were run in SDS-PAGE and the mTORC1-mediated phosphorylation on 4E-BP1^T37/46^ was detected by immunoblotting with a specific antibody (#9459, Cell Signalling Technology).

### Immunofluorescence and confocal microscopy

Immunofluorescence/confocal microscopy experiments were performed as described previously ^24,100^. In brief, cells were seeded on glass coverslips (coated with fibronectin), treated as described in the figure legends, and fixed with 4% paraformaldehyde (PFA) in 1x PBS (10 min, room temperature), followed by two permeabilization/washing steps with PBT (1x PBS, 0.1 % Tween-20). Cells were blocked in BBT (1x PBS, 0.1% Tween-20, 1% BSA) for 45 minutes. Staining with anti-mTOR (#2983, Cell Signaling Technology), anti-RagC (#9480, Cell Signaling Technology), and anti-LAMP2 (#H4B4, Developmental Studies Hybridoma Bank) primary antibodies diluted 1:200 in BBT solution was performed for 2h at room temperature. Staining with anti-TFEB (#4240, Cell Signaling Technology) or anti-TFE3 (#14779, Cell Signaling Technology) antibodies (1:200 in BBT) was performed by incubation for 16 h at 4°C. After staining with primary antibodies, cells were washed three times with PBT. Next, cells were stained with highly cross-adsorbed fluorescent secondary antibodies (Donkey anti-rabbit Alexa Fluor 488, Donkey anti-mouse TRITC; both from Jackson ImmunoResearch) diluted 1:200 in BBT for 1 hour. Nuclei were stained with DAPI (#A1001, VWR) (1:2000 in PBT) for 5 min and coverslips were washed three times with PBT solution before mounting on glass slides with Fluoromount-G (#00-4958-02, Invitrogen).

For LC3B or p62 staining, cells were fixed with 100% methanol for 15 minutes at −20°C, permeabilized with 0,1% Triton X-100 (#A4975, AppliChem) for 5 minutes and blocked for 1 hour in LC3B blocking solution (1x PBS, 5% FBS, 0,3% Triton X-100). Coverslips were incubated overnight at 4 °C with anti-LC3B (#3868, Cell Signaling Technology) or anti-p62 (#5114, Cell Signaling Technology) antibodies in LC3B/p62 staining solution (1x PBS, 1% BSA, 0,3% Triton X-100). Slides were washed three times in 1x PBS, incubated with Donkey anti-rabbit Alexa Fluor 488 (Jackson ImmunoResearch) (1:500, in 1x PBS, 1% BSA, 0,3% Triton X-100) for 1 hour at room temperature. Coverslips were then washed twice with 1x PBS, stained with DAPI (1:2000 in 1x PBS) and mounted on glass slides with Fluoromount-G (#00-4958-02, Invitrogen). All images were captured on an SP8 Leica confocal microscope (TCS SP8 X or TCS SP8 DLS, Leica Microsystems) using a 40x oil objective lens. Image acquisition was performed using the LAS X software (Leica Microsystems). Images from single channels are shown in grayscale, whereas in merged images, Alexa Fluor 488 is shown in green, TRITC in red and DAPI in blue.

### LysoTracker staining

For LysoTracker staining experiments, cells were seeded in fibronectin-coated coverslips and grown until they reached 80-90% confluency. Lysosomes were stained by the addition of 100 nM LysoTracker Red DND-99 (#L7528, Invitrogen) in complete media for 1 hour in standard culturing conditions. Cells were then fixed with 4% PFA in PBS for 10 min at room temperature, washed and permeabilized with PBT solution (1x PBS, 0.1% Tween-20), and nuclei stained with DAPI (1:2000 in PBT) for 10 min. Coverslips were mounted on slides using Fluoromount-G (#00-4958-02, Invitrogen). All images were captured on an SP8 Leica confocal microscope (TCS SP8 X or TCS SP8 DLS, Leica Microsystems) using a 40x oil objective lens. Image acquisition was performed using the LAS X software (Leica Microsystems).

### Quantification of colocalization

Colocalization analysis in confocal microscopy experiments was performed as in ^39,100^, using the Coloc2 plugin of the Fiji software ^101^. An average of 50 individual cells from 3-5 independent representative images per condition captured from one representative experiment (out of 2-3 independent replicate experiments as indicated in the figure legends) was used to calculate Manders’ colocalization coefficient (MCC) with automatic Costes thresholding ^102–104^. For experiments in which lysosomal size and morphology are affected (for instance in BafA1-treated or GNPTAB KO cells), thus also influencing lysosomal signal distribution or intensity, Pearson’s correlation coefficient (PCC) was used instead ^104^. The area corresponding to the cell nucleus was excluded from the cell region of interest (ROI) to prevent false-positive colocalization due to automatic signal adjustments. MCC and PCC are defined as a part of the signal of interest (mTOR or RagC), which overlaps with a second signal (LAMP2).

### Quantification of LC3B, p62, and LysoTracker intensities

Signal intensity was calculated using the Fiji software. Regions-of-interest (ROIs) were determined for approximately 50 cells over 5 independent representative images per condition and integrated density was calculated, representing the sum of the values of all pixels in the given ROI. Exact numbers of individual cells analysed per experiment are indicated in the figure legends.

### Scoring of TFEB/TFE3 localization

Subcellular localization of TFEB and TFE3 was performed by scoring the distribution of signal in the cytoplasm and the nucleus. Five independent fields per condition were analysed for each experiment. Exact numbers of individual cells analysed per experiment are indicated in the figure legends.

### OPP assay

To test *de novo* protein synthesis, OPP (O-propargyl-puromycin) incorporation assays were performed using the Click-iT Plus OPP Protein Synthesis Assay kit (#C10456, Thermo Fisher Scientific), according to the manufacturer’s instructions. In brief, cells seeded in fibronectin-coated coverslips until they reached 80-90% confluence. Control samples were treated with 100 µM cycloheximide (#239765, Sigma) for 4 hours before fixation to block translation. Click-iT OPP component A (20 µM) was added to the culture media for 30 minutes, cells were fixed for 10 min at room temperature with 4% PFA, and washed twice with PBT. Next, cells were incubated with Click-iT Plus OPP reaction cocktail for 30 minutes at room temperature protected from light, followed by one wash with Click-iT Reaction Rinse Buffer and further DAPI staining as described for immunofluorescence. All samples were imaged on an SP8 Leica confocal microscope (TCS SP8 X or TCS SP8 DLS, Leica Microsystems) using a 40x oil objective lens. Image acquisition was performed using the LAS X software (Leica Microsystems).

For cytometry-based detection of protein translation levels, 1×10^6^ cells were used per condition. Cells were incubated with 20 µM of Click-iT OPP component A for 1 hour, harvested and centrifuged for 3 min, 1400 rpm at room temperature. Samples were fixed with ice-cold 70% ethanol, incubated for 30 min on ice, washed once with 1x PBS, followed by two washes with 1x PBS, 0.3% BSA, and permeabilization with 0.1% Saponin in 1x PBS, 0.3% BSA for 10 min at room temperature. Cells were then centrifuged for 3 min, 1400 rpm at room temperature, the supernatant was removed and cells were incubated with Click-iT Plus OPP reaction cocktail for 30 minutes. Next, cells were washed twice with 1x PBS, 0.3% BSA and Alexa Fluor 488 signal was detected with a FITC filter in a BD LSR Fortessa Cell Analyzer flow cytometer (BD Biosciences) and further analysed in the FlowJo v10 software (TreeStar).

### Gene expression analysis

To assess mRNA expression of selected SREBP, HIF1, ATF4, and TFEB target genes, we used two RNA-seq datasets: from rapamycin- or DMSO-treated HEK293FT cells (described in ^59^; NCBI Sequence Read Archive PRJNA872474), and from RagA/B KO versus control HEK293FT cells. Target genes of each transcription factor were retrieved using the Harmonizome 3.0 online platform (https://maayanlab.cloud/Harmonizome/) ^105^. For ATF4 targets, a list of non-redundant genes was assembled using the ATF4 gene set from Harmonizome 3.0 and previous literature ^106,107^. Genes for follow-up analyses were selected following manual validation in the literature. Dot plots showing the changes in the expression of the selected target genes in each of the two RNA-seq datasets for each transcription factor were generated using the Scatter Plot tool of the Flaski toolbox ^108^ (https://flaski.age.mpg.de, developed and provided by the MPI-AGE Bioinformatics core facility). For each dot, colour and size indicate log_2_-transformed fold change (Log_2_FC), and outline colour indicates significance (black: adj. p-value < 0.05; grey: adj. p-value ≥ 0.05).

### Statistical analysis

Statistical analysis and presentation of quantification data was performed using GraphPad Prism (versions 9.1.0 and 9.2.0). Data in graphs in Fig. 5g and Extended Data Fig. 4a and 7b are shown as mean ± SD. Data in all other graphs are shown as mean ± SEM. For graphs with only two conditions shown, significance for pairwise comparisons was calculated using Student’s t-test. For graphs with three or more conditions shown, significance for pairwise comparisons to the respective controls was calculated using one-way ANOVA with *post hoc* Holm-Sidak test. Sample sizes (n) and significance values are indicated in figure legends (* p < 0.05, ** p < 0.01, *** p < 0.001, **** p < 0.0001, ns: non-significant).

All findings were reproducible over multiple independent experiments, within a reasonable degree of variability between replicates. The number of replicate experiments for each assay is provided in the respective figure legends. No statistical method was used to predetermine sample size, which was determined in accordance with standard practices in the field. No data were excluded from the analyses. The experiments were not randomized, and the investigators were not blinded to allocation during experiments and outcome assessment.

## Acknowledgements

We thank all members of the Demetriades lab for critical discussions; Andreas Lamprakis for technical support; the MPI-AGE FACS & Imaging Core Facility for support with confocal microscopy and FACS analysis. SAF, JP and FA received support by the Cologne Graduate School of Ageing Research. CD is funded by the European Research Council (ERC) under the European Union’s Horizon 2020 research and innovation programme (grant agreement No 757729), and by the Max Planck Society. Parts of this work were supported by the Deutsche Forschungsgemeinschaft (DFG, German Research Foundation) through the Research Unit Grant FOR2722 (Project No 384170921) to CD. Figures created with BioRender.com.

## Author Contributions

Experimental work: SAF, DDA, JN, JP, PG, YE, SW, MKS; data analysis: SAF, DDA, FA, CD; project design & conceptualisation: CD; project supervision: CD; funding acquisition: CD; figure preparation: SAF, DDA, FA, CD; manuscript draft: CD, with contributions from all authors. All authors approved the final version of the manuscript and agree on the content and conclusions.

## Declaration of interests

The authors declare no competing interests.

## Data availability

The data that support the findings of this study (uncropped immunoblots, microscopy pictures) are available from the corresponding author upon reasonable request.

## Code availability

No code was generated in this study.

## Additional Information

Supplementary Information (Extended Data Figures 1-8 and Supplementary Tables 1-2) is available for this paper.

Correspondence and requests for materials should be addressed to Constantinos Demetriades.

## Notes

### Competing Interest Statement

The authors have declared no competing interest.

