## Supplementary material for "Spatially and Functionally Distinct mTORC1 Entities Orchestrate the Cellular Response to Amino Acid Availability": Suppl Figures 1-8

### Supplementary Information

Supplementary Tables 1-2 and Extended Data Figures 1-8

#### Supplementary Tables

Supplementary Table 1. List of primary antibodies used in this study.

Supplementary Table 2. List of DNA oligonucleotides used in this study.

#### Extended Data Figures

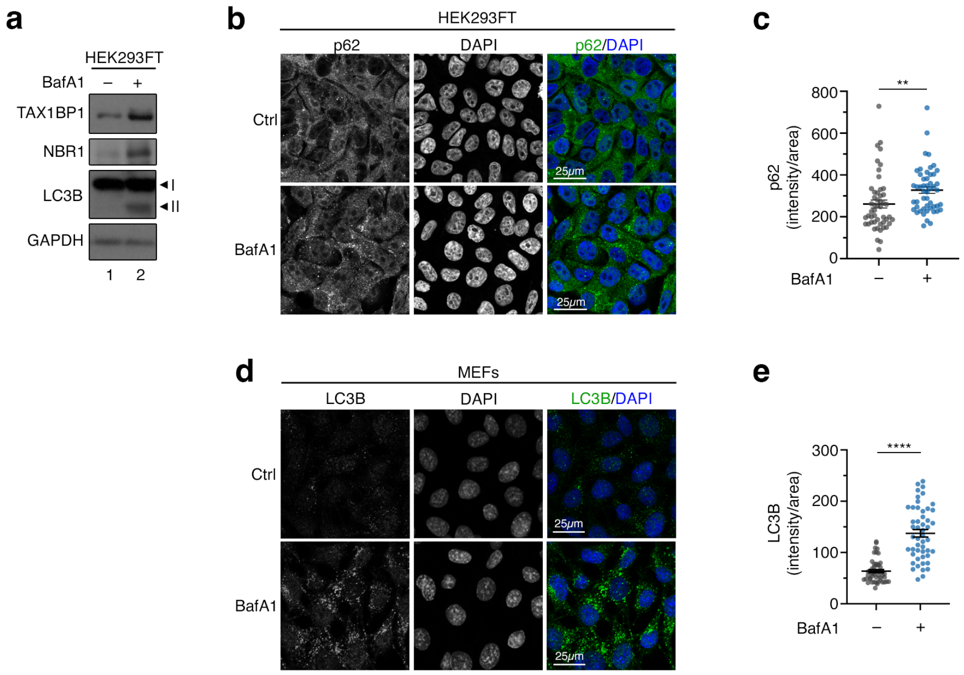


**Extended Data Figure 1. Basal lysosomal degradative activity takes place also in cells grown in the presence of exogenous nutrients.**

**(a)** Immunoblots with lysates from HEK293FT WT cells, treated with BafA1 as shown, probed with the indicated antibodies. n = 3 independent experiments

**(b-c)** p62 accumulates in BafA1-treated cells, shown by immunofluorescence and confocal microscopy. Scale bars = 25 μm (b). Quantification of p62 signal from n = 50 individual cells from 5 independent fields per condition in (c). Representative data from one out of two independent replicate experiments are shown.

**(d-e)** Basal lysosomal degradative activity in MEFs indicated by accumulation of LC3B upon BafA1 treatment. Scale bars = 25 μm (d). Quantification of LC3B signal from n = 48-50 individual cells from 5 independent fields per condition in (e). Representative data from one out of three independent replicate experiments are shown.

Data in graphs shown as mean ± SEM. ** p<0.01, **** p<0.0001.

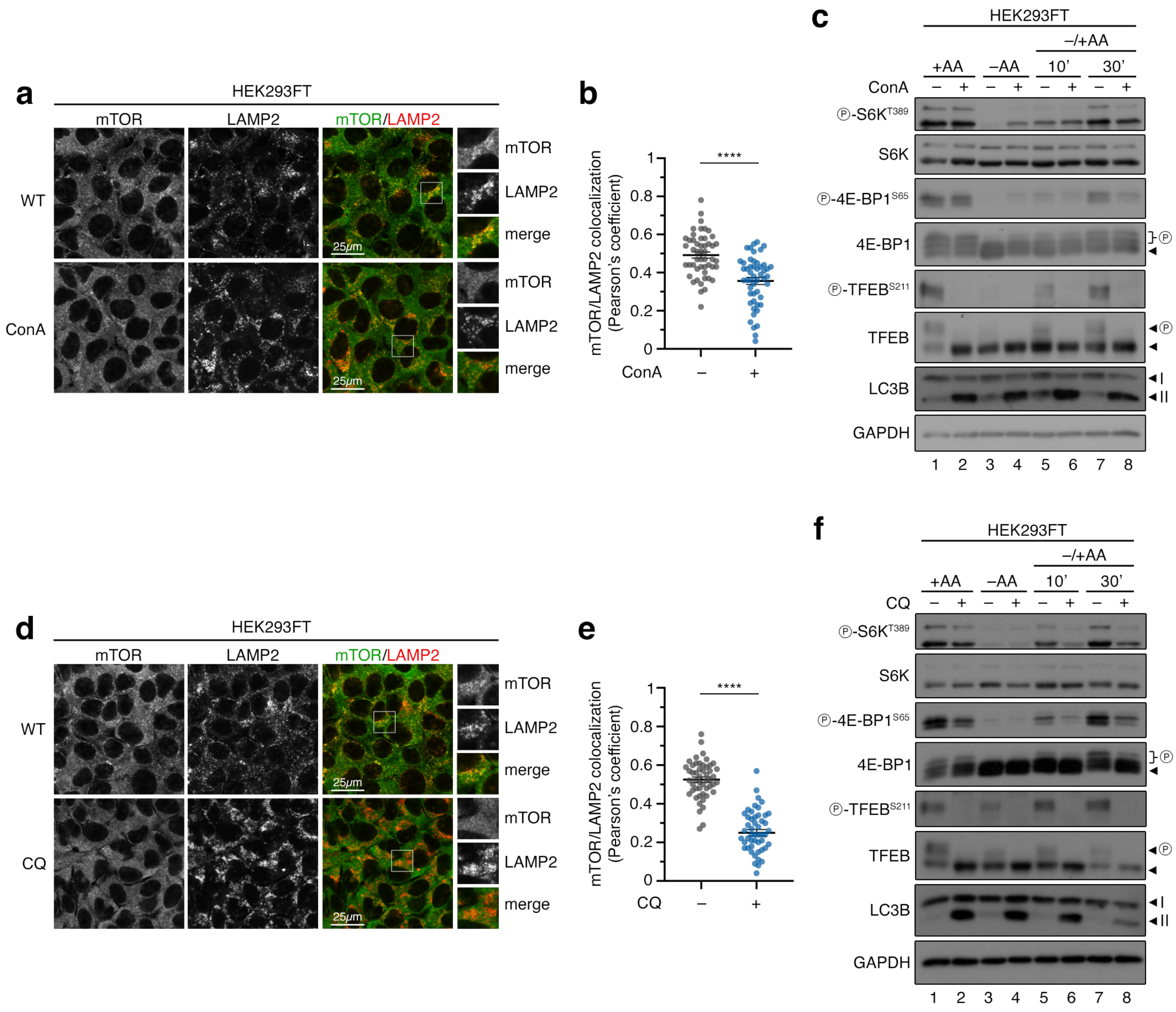


**Extended Data Figure 2. Blockage of lysosomal function using ConA or CQ disconnects mTORC1 localization on lysosomes and activity toward cytoplasmic substrates.**

**(a-b)** Lysosomal accumulations of mTOR are lost in concanamycin A (ConA)-treated cells. Scale bars = 25 μm (a). Quantification of mTOR/LAMP2 colocalization in (b). n = 50 individual cells from 5 independent fields per condition.

**(c)** ConA treatment preferentially diminishes phosphorylation of the lysosomal substrate TFEB but not of the cytoplasmic substrates S6K and 4E-BP1 under basal culture conditions. n = 3 independent experiments.

**(d-f)** As in (a-c) but for treatments with chloroquine (CQ).

Arrowheads indicate bands corresponding to different protein forms, when multiple bands are present. P: phosphorylated form. Representative data from one out of three independent replicate experiments are shown. Data in graphs shown as mean ± SEM. **** $p<0.0001$.

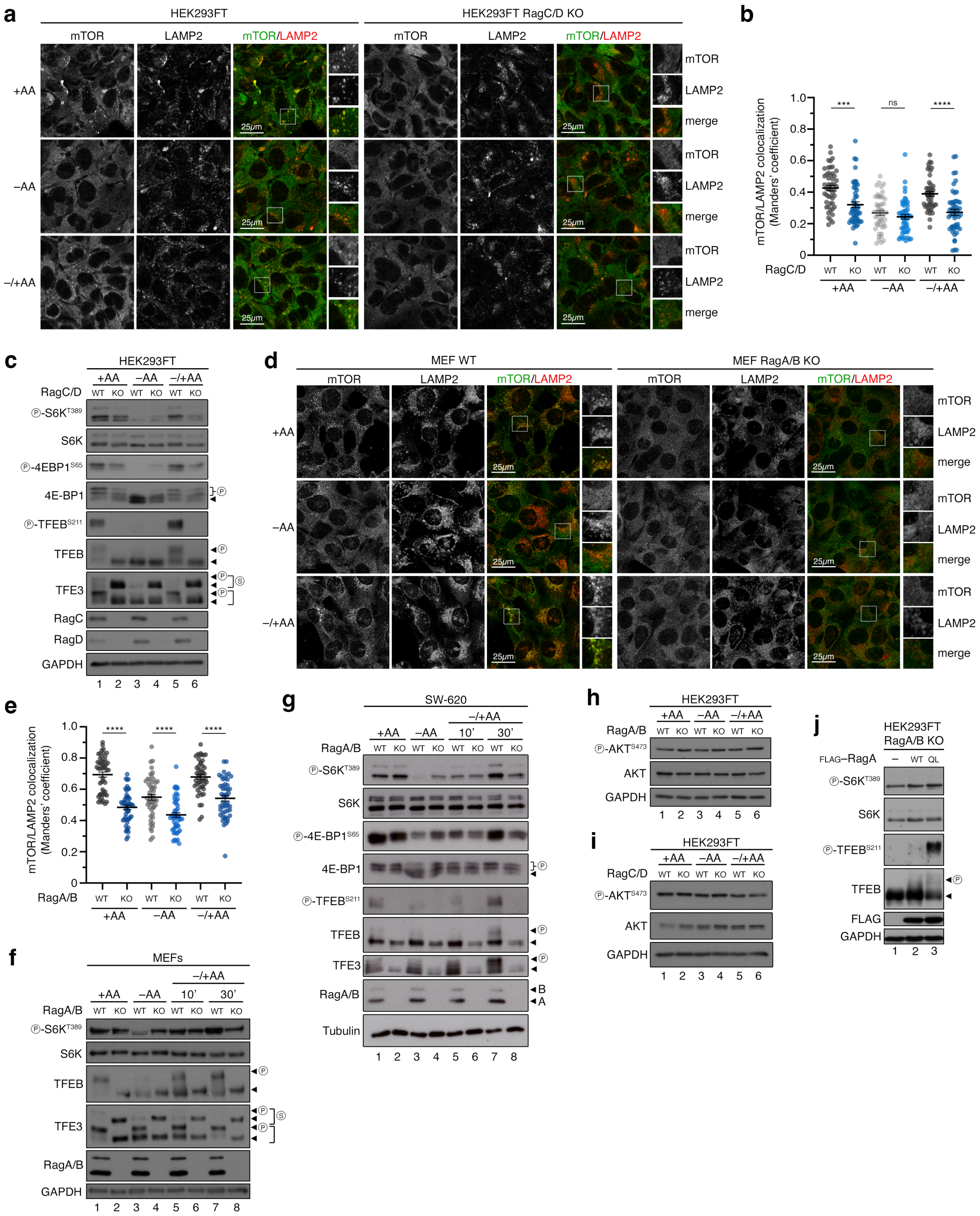


**Extended Data Figure 3. Rag loss-of-function disconnects mTORC1 localization and activity in different cell lines, without affecting mTORC2, and is reversible by RagA re-expression.**

**(a-b)** Colocalization analysis of mTOR with LAMP2 (lysosomal marker) in HEK293FT WT or RagC/D KO cells, treated as indicated in the figure, using confocal microscopy. Scale bars = 25 μm. Magnified insets shown to the right (a). Quantification of colocalization in (b). n = 50 individual cells from 5 independent fields per condition. Representative data from one out of three independent experiments are shown.

**(c)** Immunoblots with lysates from HEK293FT WT and RagC/D KO cells, treated with media containing or lacking AA, in basal (+AA), starvation (–AA) or add-back (–/+AA) conditions, probed with the indicated antibodies. n = 3 independent experiments.

**(d-e)** Colocalization analysis of mTOR with LAMP2 (lysosomal marker) in MEF WT or RagA/B KO cells, treated as indicated in the figure, using confocal microscopy. Scale bars = 25 μm. Magnified insets shown to the right (d). Quantification of colocalization in (e). n = 50 individual cells from 3 independent fields per condition.

**(f)** As in (c), but with WT and RagA/B KO MEFs.

**(g)** As in (c), but with WT and RagA/B KO SW-620 cells. n = independent experiments.

**(h)** Immunoblots with lysates from HEK293FT WT and RagA/B KO cells, treated with media containing or lacking AA, in basal (+AA), starvation (–AA) or add-back (–/+AA) conditions, probed with the indicated antibodies.

**(i)** As in (h), but with WT and RagC/D KO HEK293FT cells. n = 3 independent experiments.

**(j)** Re-expression of WT or active-locked RagA (QL) in RagA/B KO cells rescues TFEB phosphorylation, without strongly affecting p-S6K levels.

Arrowheads indicate bands corresponding to different protein forms, when multiple bands are present. P: phosphorylated form; S: SUMOylated form. Data in graphs shown as mean ± SEM. *** p<0.001, **** p<0.0001, ns: non-significant.

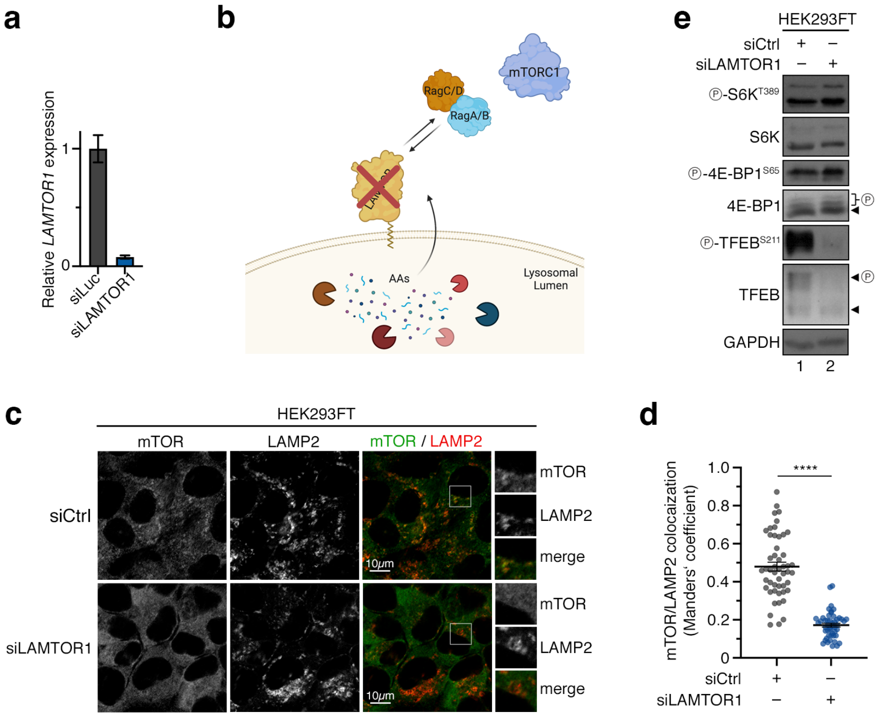


**Extended Data Figure 4. LAMTOR1/p14 knockdown recapitulates Rag loss-of-function.**

**(a)** Expression analysis of *LAMTOR1* by qPCR confirms successful knockdown in HEK293FT cells. Data shown as mean ± SD.

**(b)** Schematic model of lysosomal tethering of the Rag dimer by the LAMTOR complex.

**(c-d)** Colocalization analysis of mTOR with LAMP2 (lysosomal marker) in HEK293FT WT cells, using confocal microscopy. Cells were transiently transfected with siRNAs targeting LAMTOR1 or a control RNAi duplex (siCtrl). Scale bars = 10 μm. Magnified insets shown to the right (c). Quantification of colocalization in (d). n = 50 individual cells from 3 independent fields per condition. Data shown as mean ± SEM. **** $p<0.001$.

**(e)** Immunoblots with lysates from HEK293FT WT cells, transiently transfected with siRNAs targeting LAMTOR1 or a control RNAi duplex (siCtrl), cultured under basal conditions, and probed with the indicated antibodies. Arrowheads indicate bands corresponding to different protein forms, when multiple bands are present. P: phosphorylated form.

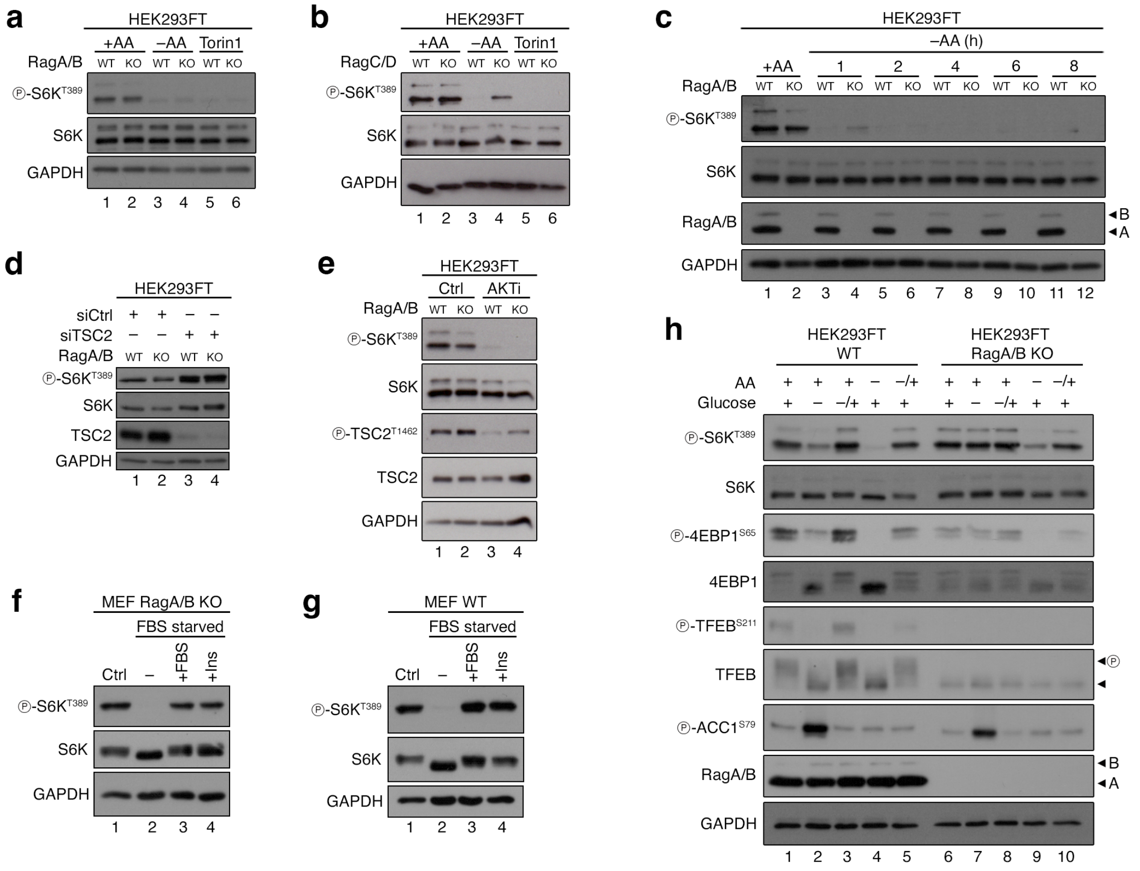


**Extended Data Figure 5. Phosphorylation of S6K by non-lysosomal mTORC1 responds to exogenous AAs and is regulated downstream of mTOR and growth factor signalling.**

**(a)** Immunoblots with lysates from WT and RagA/B KO HEK293FT cells, treated with media containing (+AA) or lacking AA (–AA), or with Torin1 as shown, probed with the indicated antibodies. n = 3 independent expriments.

**(b)** As in (a), but for WT and RagC/D HEKs. n = 2 independent experiments.

**(c)** AA starvation time-course in WT and RagA/B KO HEK293FT cells shows that the residual S6K phosphorylation in the KOs at the early time points of starvation disappears at slightly later times. n = 3 independent experiments.

**(d)** Immunoblots with lysates from WT and RagA/B KO HEK293FT cells, transiently transfected with siRNAs targeting TSC2 or a control RNAi duplex (siCtrl), probed with the indicated antibodies.

**(e)** Immunoblots with lysates from WT and RagA/B KO HEK293FT cells, treated with Akt inhibitor (AKTi) as shown, probed with the indicated antibodies. n = 3 independent experiments.

**(f-g)** Growth factor removal and re-addition is sensed similarly in RagA/B KO (f) and WT MEFs (g). Cells were starved for 1 h in media lacking FBS (–) and then re-stimulated with FBS (+FBS; final concentration 10%) or insulin (+Ins) for 30 min.

**(h)** Rag KO cells are deficient in sensing glucose depletion, in line with previous studies. WT and RagA/B KO HEKs were cultured under basal glucose- and AA-replete conditions, or starved with media lacking glucose or AAs for 1h (–), or first starved and then re-stimulated (–/+) with glucose or AAs for 30 min. n = 2 independent experiments.

Arrowheads indicate bands corresponding to different protein forms, when multiple bands are present. P: phosphorylated form.

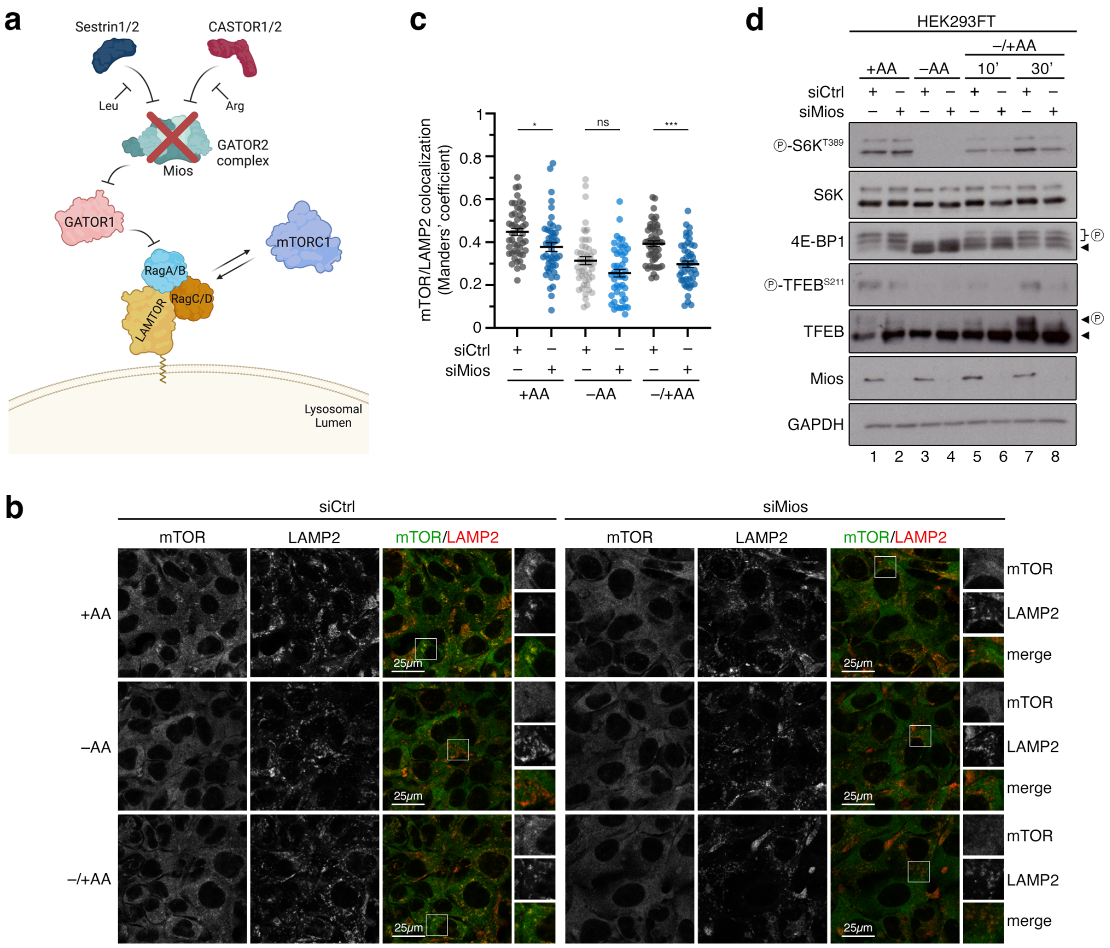


**Extended Data Figure 6. Non-lysosomal mTORC1 is regulated independently from the GATOR2 complex.**

**(a)** Schematic model of cytoplasmic AA sensing and signaling upstream of the Rags. See text for details.

**(b-c)** Colocalization analysis of mTOR with LAMP2 (lysosomal marker) in HEK293FT WT cells, using confocal microscopy. Cells were transiently transfected with siRNAs targeting Mios or a control RNAi duplex (siCtrl) and treated as indicated. Scale bars = 25 μm. Magnified insets shown to the right (b). Quantification of colocalization in (c). n = 46-50 individual cells from 5 independent fields per condition. Representative data from one out of two independent experiments are shown.

**(d)** Immunoblots with lysates from HEK293FT WT cells, transiently transfected with siRNAs targeting Mios or a control RNAi duplex (siCtrl), and treated with media containing or lacking AA, in basal (+AA), starvation (–AA) or add-back (–/+AA) conditions, probed with the indicated antibodies. n = 3 independent experiments.

Arrowheads indicate bands corresponding to different protein forms, when multiple bands are present. P: phosphorylated form. Data in (c) shown as mean ± SEM. * $p<0.05$, *** $p<0.001$, ns: non-significant.

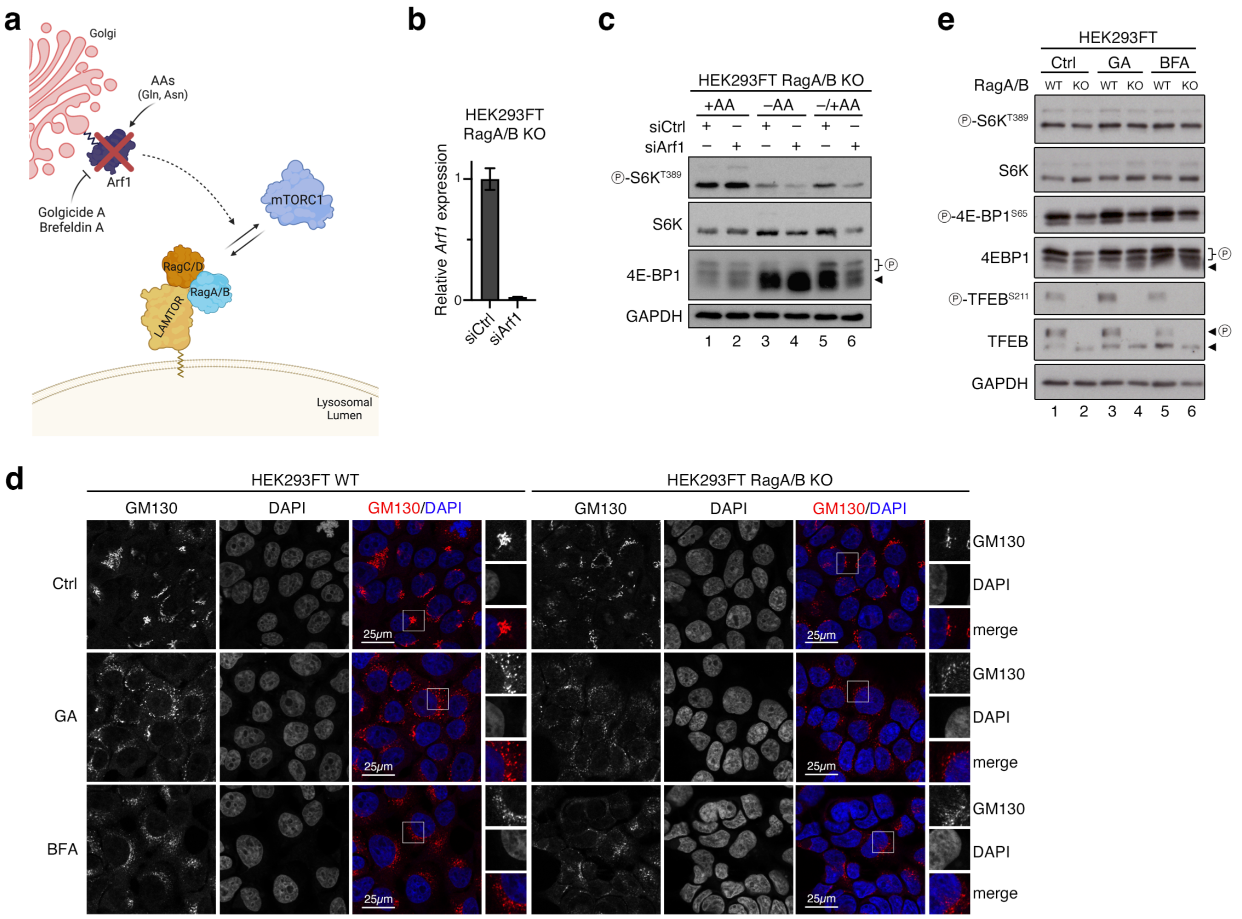


**Extended Data Figure 7. Non-lysosomal mTORC1 activity is independent from Arf1 and the Golgi apparatus.**

**(a)** Schematic model of Golgi-localized Arf1 regulating mTORC1 localization and activity in response to re-addition of specific AAs. Golgicide A (GA) or Brefeldin A (BFA) block Arf1 via targeting the ArfGEF GBF1.

**(b)** Expression analysis of *Arf1* by qPCR confirms successful knockdown in RagA/B KO HEK293FT cells. Data shown as mean ± SD.

**(c)** Immunoblots with lysates from RagA/B KO HEK293FT cells, transiently transfected with siRNAs targeting Arf1 or a control RNAi duplex (siCtrl), and treated with media containing or lacking AAs, in basal (+AA), starvation (–AA) or add-back (–/+AA) conditions, probed with the indicated antibodies.

**(d)** Golgi morphology using GM130 (Golgi marker) in HEK293FT WT and RagA/B KO cells, using confocal microscopy. Cells treated with Golgicide A (GA) or Brefeldin A (BFA) as indicated. Magnified insets shown to the right. Scale bars = 25 μm. n = 3 independent experiments.

**(e)** Immunoblots with lysates from WT and RagA/B KO HEK293FT cells, treated with GA or BFA as shown in the panel, probed with the indicated antibodies. n = 3 independent experiments. Arrowheads indicate bands corresponding to different protein forms, when multiple bands are present. P: phosphorylated form.

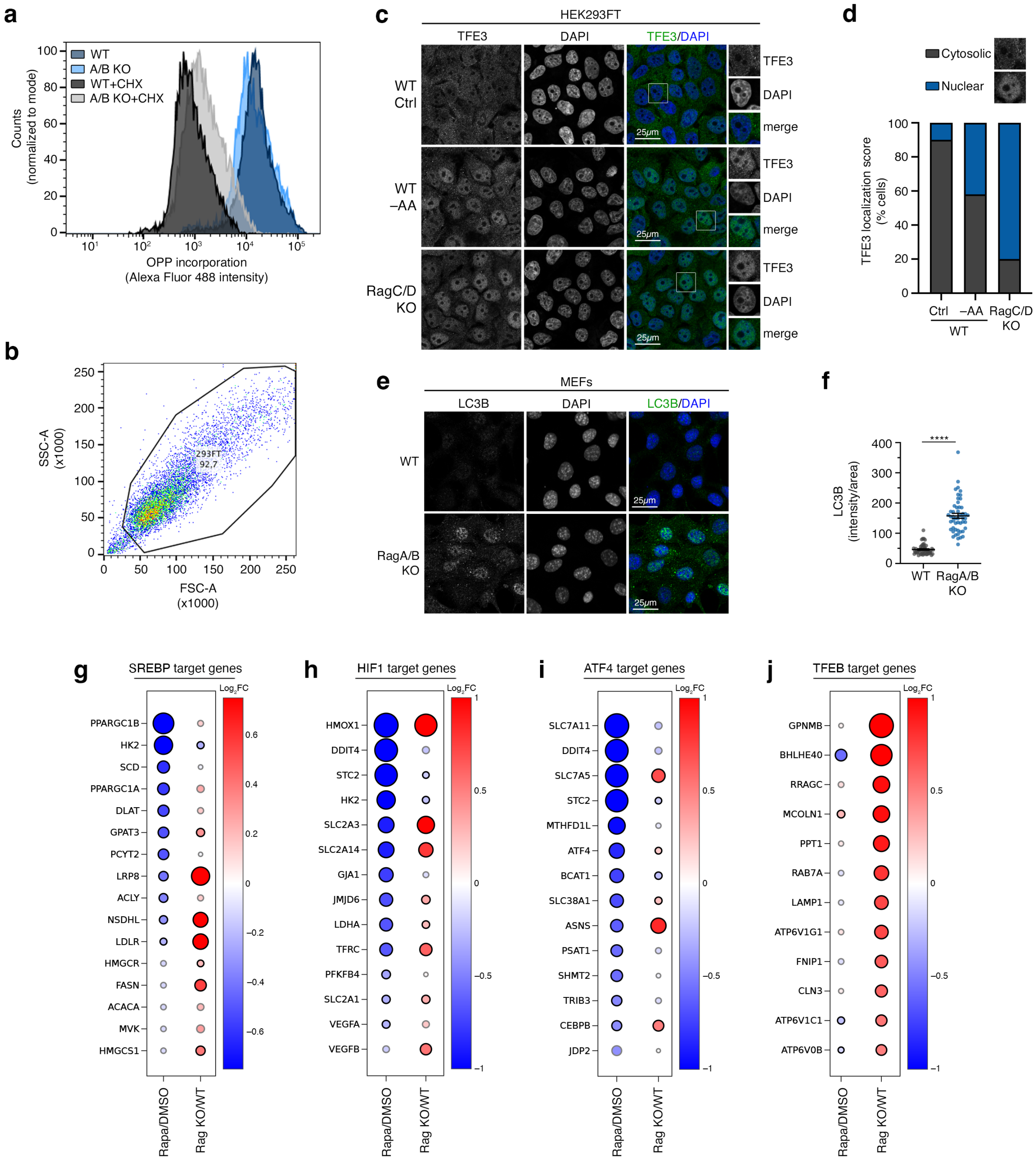


**Extended Data Figure 8. Functional separation of Rag-dependent and -independent mTORC1 activities.**

**(a)** *De novo* protein synthesis (OPP incorporation) assay with WT and RagA/B KO HEK293FT cells, analysed by flow cytometry. Cycloheximide (CHX) used as control to block protein synthesis. Cell counts: $n_{WT}$ = 9306, $n_{ABKO}$ = 9317, $n_{WT+CHX}$ = 9572, $n_{ABKO+CHX}$ = 8869. Representative data from one out of three independent experiments are shown.

**(b)** Gating strategy for the flow-cytometry-based OPP incorporation assay in (a).

**(c-d)** TFE3 localization analysis in WT and RagC/D KO HEK293FT cells, using confocal microscopy. AA starvation (–AA) was used as control. Nuclei stained with DAPI. Magnified insets shown to the right. Scale bars = 25 μm (c). Scoring of TFE3 localization in (d). Individual cells were scored for nuclear or cytoplasmic TFEB localization as indicated in the example images. $n_{WT\_Ctrl}$ = 81 cells, $n_{WT\_-AA}$ = 92 cells, $n_{ABKO}$ = 103 cells. Representative data from one out of three independent experiments are shown.

**(e-f)** LC3B staining in WT and RagA/B KO MEFs. Nuclei stained with DAPI. Scale bars = 25 μm (e). Quantification of LC3B signal (50 individual cells from 5 independent fields per condition) in (f). n = 2 independent experiments.

**(g)** Gene expression analyses from RNA-seq experiments comparing rapamycin- to control-treated HEK293FT cells (Rapa/DMSO) or RagA/B KO to WT HEK293FT cells (Rag KO/WT) indicate SREBP activity is not consistently affected by loss of the Rags. Dot plot showing the changes in the expression of selected SREBP target genes in each of the two datasets. For each dot, colour and size indicate $log_2$-transformed fold change ($Log_2FC$), and outline colour indicates significance (black: adj. p-value < 0.05; grey: adj. p-value ≥ 0.05).

**(h)** As in (g), but for selected HIF1 target genes.

**(i)** As in (g), but for selected ATF4 target genes.

**(j)** As in (g), but for selected TFEB target genes. Note the specific upregulation of lysosome-related genes in the RagA/B KO vs. WT comparison, whereas no such changes are observed in the respective Rapamycin vs. DMSO comparison.

Data in (f) shown as mean ± SEM. **** p<0.0001.
